# Florigen Activation Complex Dynamics and SVP-Mediated Repression Orchestrate Temperature-Regulated Flowering in Saffron

**DOI:** 10.1101/2025.07.01.662559

**Authors:** Diksha Kalia, Joel Jose-Santhi, Firdous Rasool Sheikh, Rajesh Kumar Singh

## Abstract

Saffron, a high-value spice cultivated worldwide for its therapeutic and culinary uses, is a sterile triploid species, rendering conventional breeding approaches ineffective. This limitation underscores the need for molecular and biotechnological strategies for its genetic improvement. Flowering, a key determinant of saffron yield, is strongly influenced by temperature; however, the genetic regulatory networks underlying this process remain poorly understood. Our study identifies key regulators of saffron’s flowering, focusing on the Florigen Activation Complex (FAC) components: *FLOWERING LOCUS T* (FT), bZIP transcription factor FD, and *TERMINAL FLOWER*-1 (TFL-1), and demonstrate their temperature-dependent roles in floral regulation. Spatiotemporal expression analyses suggested that *CsatFT3* and *CsatFD2*, expressed in the floral meristem promote floral induction, while *CsatTFL1*-3 acts as a floral repressor. Protein interaction studies showed that CsatFT3 and CsatTFL1-3 compete for binding to CsatFD2, and their balance modulates floral induction. Functional validation in Arabidopsis and Saffron confirmed these findings. Furthermore, we identified *CsatSVP2*, an ortholog of *SHORT VEGETATIVE PHASE* (SVP), as a low temperature-responsive repressor that directly binds the CsatFT3 promoter to inhibit its expression. Together, these findings enhance our understanding of temperature mediated floral induction in saffron and provide insights and lay the groundwork for genetic interventions to enhance yield under variable temperature conditions.

## Introduction

Saffron (*Crocus sativus* L.) is a sterile, clonally propagated, autotriploid geophytic monocot cultivated for its highly valued dried stigmas, which constitute the world’s most expensive spice. While the stigmas are the primary edible and culinary component, other parts of the saffron plant also have diverse applications, including uses in agriculture, traditional medicine, and as a natural colouring agent, making the entire flower economically valuable (Ahrazem et al., 2015). Saffron has been predominantly cultivated for centuries in select regions such as Iran, India, Greece, Italy, Afghanistan, Morocco, Spain, and various Mediterranean basins (Cardone et al., 2020). Flowering is the key determinant of saffron’s crop productivity, and this critical phase is increasingly influenced by environmental changes. As a thermoperiodic plant, saffron’s flowering is primarily regulated by temperature. During the warm summer months, the transition from vegetative to reproductive growth occurs underground, with floral bud emergence only triggered by exposure to cooler temperatures (Molina et al., 2005a). This biphasic thermal requirement consists of an induction phase at warm temperatures lasting 50–150 days, followed by an emergence phase under cooler conditions, a pattern that has been well documented (Jose-Santhi et al., 2023; Molina et al., 2005a). The inappropriate temperature during the flowering transition leads to flower atrophy or no flower, causing yield loss (Wang et al., 2021). Saffron corms can sense the temperature change and modify the response accordingly (Molina et al., 2005b). Thus, there exists a thermoresponsive regulation of flowering in saffron which is still not well studied at the molecular level.

Flowering is a key transition in the angiosperm life cycle, marking the shift from vegetative to reproductive growth and representing a tightly regulated process vital for survival. This process is orchestrated by a complex network of environmental cues, internal signalling pathways, hormonal regulation, and genetic programs that together ensure flowering occurs under optimal conditions (Zik and Irish, 2003). Key environmental pathways influencing flowering include photoperiod, ambient temperature, light quality, vernalization, and gibberellin signalling. These pathways operate in coordination with endogenous factors along with plant age and nutritional status, allowing plants to synchronize reproduction with favourable environmental windows (Andrés and Coupland, 2012; Freytes et al., 2021; Pyo et al., 2014; Srikanth and Schmid, 2011). Extensive studies in both monocot and dicot organisms, such as Arabidopsis thaliana and rice, have uncovered key regulatory components of flowering (Kinoshita and Richter, 2020). Central to this regulation are floral integrator genes, particularly *FLOWERING LOCUS T* (*FT*) in Arabidopsis and *Heading Date 3a* (*Hd3a*) in rice, which belong to the phosphatidylethanolamine-binding protein (PEBP) family. FT/Hd3a promotes flowering by forming the Florigen Activation Complex (FAC), composed of FD-like bZIP transcription factors and the florigen receptor 14-3-3 protein (Abe et al., 2005; Taoka et al., 2013). This complex activates floral meristem identity genes such as *LEAFY* and *APETALA1*, initiating floral development (Putterill et al., 2004; Simon et al., 1996; Wigge et al., 2005). In contrast, *TERMINAL FLOWER 1* (TFL1), another PEBP family member, acts antagonistically to repress flowering (Wickland and Hanzawa, 2015). In rice, Hd3a, RFT1, and OsFTL10 promote flowering by forming the Florigen Activation Complex (FAC) (Komiya et al., 2008; Tamaki et al., 2007). Conversely, OsFTL12, OsFTL4, RCN1, and RCN2 act as floral repressors by competing for FAC components to assemble a Florigen Repression Complex (FRC) (Nakagawa et al., 2002; Zheng et al., 2023). The dynamic balance between FAC and FRC finely regulates heading date and plant architecture in rice. Members of the FD family are key interaction partners of the PEBP family, playing roles in both promoting and repressing meristem differentiation, depending on their specific interacting partners (Zhu et al., 2020). Concurrently, the flowering program suppresses negative regulators, including *FLOWERING LOCUS C* (FLC), and *SHORT VEGETATIVE PHASE* (SVP), which are crucial for ensuring a proper transition from the vegetative to the reproductive phase mainly regulated by temperature (Bowman et al., 2012; Turck et al., 2008). Temperature-dependent flowering has been widely studied, with FT-like genes playing a central role (Capovilla et al., 2015). Upstream regulators include MADS-domain transcription factors such as SVP and FLC, which modulate flowering by forming complexes and repressing *FT* expression via direct binding to its promoter (Balasubramanian et al., 2006; Gu et al., 2013; Lee et al., 2007). However, the thermoresponsive pathway governing floral induction in saffron remains largely unexplored.

In contrast to annuals, the regulation of flowering in perennial geophytes involves modified or additional mechanisms to accommodate their extended life cycles, periods of dormancy and dual reproduction (Khosa et al., 2021). This has been observed in species like *Narcissus tazetta* (Noy-Porat et al., 2013), tulip (Tulipa spp.) (Leeggangers et al., 2017a), and saffron (Molina et al., 2005a), where floral induction occurs even in the absence of leaves. In these species, flowering is thought to be mediated directly at the meristem level, independent of traditional photoperiod or vernalization cues. Instead, ambient temperature plays a dominant role in determining flowering time. These observations highlight the need to identify the specific genes and mechanisms that govern flowering in geophytic perennials. Over evolutionary time, the molecular mechanisms controlling flowering have grown increasingly complex, involving intricate interactions between floral regulator genes and their transcription factor partners, which form regulatory complexes (Wickland and Hanzawa, 2015). Gene duplication and neofunctionalization have contributed to the diversification of these regulatory genes. Both *FT* and *TFL1,* the central regulators of flowering, have undergone gene duplication events in various lineages, leading to paralogs with distinct functions that promote or inhibit flowering (Jin et al., 2021). This evolutionary diversification is evident across many species. In geophytes, for instance, members of the PEBP gene family have extended their functions beyond flowering to include the formation of underground storage organs, contributing to both sexual and asexual reproduction (Khosa et al., 2021). In species like potato, tulip, and others, gene duplication has led to distinct roles in flowering, vegetative growth, and tuber or bulb formation (Bellinazzo et al., 2025; Jing et al., 2023; Navarro et al., 2011). Similar divergence has been observed in onion (*Allium cepa*), sugar beet (*Beta vulgaris*), and hybrid aspen, demonstrating the complexity of flowering regulation in perennials (Hsu et al., 2011; Lee et al., 2013b; Pin et al., 2010). Beyond the PEBP gene family, functional diversification is seen in related pathways. In hybrid aspen, two FD homologs, *FDL1* and *FDL2*, regulate seasonal growth differently (Tylewicz et al., 2015). In rice, *OsFD2* contributes to leaf development while *OsFD1* regulates flowering (Tsuji et al., 2013). FD-like transcription factors can have dual roles, promoting or repressing flowering depending on their partners. For instance, *CmFDa* in *Chrysanthemum morifolium* represses flowering through epigenetic regulation (Xue et al., 2025). These examples highlight the evolutionary plasticity of flowering networks and the importance of species-specific regulatory mechanisms.

Saffron, due to its sterile nature and extremely limited genetic diversity, is not amenable to conventional breeding approaches, which hinders efforts to develop climate-resilient, high-yielding, and stress-tolerant varieties. As a result, there have been no significant advancements in improving saffron’s adaptability to climate change, making its cultivation increasingly vulnerable to environmental fluctuations. Consequently, only biotechnological interventions can be employed to introduce desirable traits. Despite its economic and agricultural importance, research on the molecular regulation of flowering in saffron remains limited. Most existing studies have focused on temperature effects on flowering physiology (Molina et al., 2005a; Wang et al., 2021), and while several flowering-related genes have been identified through transcriptomic and genomic analyses (Hu et al., 2020; Kalia et al., 2022; Renau-Morata et al., 2021; Singh et al., 2023; Tsaftaris et al., 2013), these remain largely descriptive and lack functional validation. Expression profiling has indicated the involvement of multiple *FT* and *TFL1* homologs in saffron flowering (Kalia et al., 2022; Renau-Morata et al., 2021; Tsaftaris et al., 2013), yet no functional characterization of these genes has been reported. To fill this gap, we present a comprehensive molecular framework for temperature-dependent floral induction in saffron, a geophytic monocot with a unique flowering phenotype. By integrating gene expression profiling, heterologous functional assays, protein–protein interaction studies, promoter binding analyses, and virus-induced gene silencing, we identify key regulatory components that mediate the floral transition in response to ambient temperature. Overall, these findings advance our understanding of temperature-mediated flowering in saffron and provide a foundation for genetic interventions aimed at improving yield under fluctuating environmental conditions

## Methods

### Plant materials and growth conditions

Saffron corms *(Crocus sativus* L.) were grown under controlled conditions at CSIR-IHBT, and samples were collected monthly from January to September, covering both the vegetative growth and flowering phases (Jose-Santhi et al., 2023; Kalia et al., 2022). To analyze gene expression patterns, apical buds, axillary buds, and corm tissues were harvested across this period, with three biological replicates per time point, each comprising pooled tissue from ten individual corms. For RT-qPCR analysis, apical buds, axillary bud, corm tissue collected during the vegetative and flowering stages were used, with detailed methods and results provided in the Supplementary Information.

ViGS experiments were conducted using corms grown under the same controlled conditions as described earlier. Corms with high flowering competency (weighing >10 g) were selected for the silencing of *CsatFT3* and *CsatFD2*. To investigate the role of *CsatSVP2*, two experimental conditions were used: (i) standard growth conditions, and (ii) a temperature-sensitive treatment in which corms were incubated at 8 °C during the flowering phase to assess temperature-dependent gene regulation. For silencing of *CsatTFL1-3/CEN1*, corms with lower flowering competency (weighing 7–9 g) were utilized.

*Arabidopsis thaliana* (Col-0) was used as the wild-type background for transformation experiments, and the ft1 mutant in the Ler background was used for functional complementation of CsatFT3. *Nicotiana benthamiana* was used for transient expression assays. Arabidopsis seeds were surface-sterilized with 4% sodium hypochlorite and germinated on half-strength Murashige and Skoog (MS) medium. To synchronize germination, seeds were stratified at 4°C for two days before being transferred to a soil mixture of peat, vermiculite, and perlite (2:1:1). All plants were cultivated in controlled growth chambers under long-day conditions (16 h light/8 h dark) at 22°C for Arabidopsis and 25°C for *Nicotiana benthamiana*.

### RNA isolation and expression analysis by RT-qPCR

Total RNA was isolated from saffron apical buds, axillary buds, corm tissues, and Arabidopsis using the Plant Total RNA Isolation Kit (Sigma-Aldrich), according to the manufacturer’s protocol. For each sample, 1 µg of total RNA was used to synthesize cDNA. Reverse transcription was performed using the RevertAid cDNA Synthesis Kit. The transcript level of each gene was quantified by quantitative real-time PCR (q-PCR) using gene-specific primers (Supplementary Table 1), generated by SnapGene Viewer 5.3.1 software (version 0.4.0) using the System (BioRad CFX Opus Real time PCR). Each expression analysis included three biological replicates, with each replicate derived from pooled tissue of ten individual corms. RT-qPCR results are presented as the mean ± standard error (SEM) of three technical replicates per biological sample.

### Sequence alignments and phylogenetic analysis

Protein sequences were aligned using the ClustalW algorithm with default parameters to generate high-quality multiple sequence alignments. Visualization of the alignments was performed using ESPript 3.x (Robert & Gouet, 2014) to highlight conserved regions and structural features. The aligned sequences were exported in Newick (.nwk) format for phylogenetic analysis. Phylogenetic tree construction was carried out using the “build” function of ETE3 v3.1.3 (Huerta-Cepas et al., 2016), implemented via the GenomeNet platform (https://www.genome.jp/tools/ete/). FastTree v2.1.8 (Price et al., 2010) was used with default settings to infer approximately maximum-likelihood phylogenetic trees.

### Plant transformation

To generate Arabidopsis transgenic lines, full-length coding sequences (CDS) of the target genes were amplified and first cloned into the pENTR entry vector using Gateway® cloning technology. Subsequently, LR recombination was performed to transfer the CDS into the destination vectors pK2GW7 (for overexpression analysis), under the control of the Cauliflower Mosaic Virus (CaMV) 35S promoter. The resulting constructs included 35S::*CsatFD1*, 35S::*CsatFD2*, 35S::*CsatFD3*, 35S::*CsatFT3*, 35S::*CsatTFL1-1*, 35S::*CsatTFL1-2*, 35S::*CsatTFL1-3*, 35S::*CsatSVP1*, and 35S::*CsatSVP2*. Primer sequences used for vector construction are provided in the supplementary table 1 file. The final recombinant plasmids were introduced into *Arabidopsis thaliana* (Col-0) via Agrobacterium tumefaciens strain GV3101 using the floral dip transformation method. Transgenic seeds were harvested individually and screened based on the appropriate selection marker. Homozygous T3 lines were identified and used for further phenotypic characterization. Flowering time was recorded as the number of days from germination to the emergence of the first floral bud.

### Yeast one hybrid (Y1H) assay

To examine the binding of CsatSVP2 to the CsatFT3 promoter, yeast one-hybrid (Y1H) assays were conducted using the Matchmaker® Gold Yeast One-Hybrid System Kit (Clontech). Promoter fragments of CsatFT3 containing the core CArG-box motif were cloned into the pAbAi vector. The resulting recombinant pAbAi-promoter plasmids were linearized with BstBI (NEB) and integrated into the genome of the Y1HGold yeast strain. Transformants were selected on synthetic dextrose (SD) medium lacking uracil (SD/-Ura). The coding sequence of CsatSVP2 was amplified and inserted into the pGADT7-AD vector. The recombinant pGADT7-AD constructs were then introduced into the Y1HGold strains containing the integrated CsatFT3 promoter. Protein–DNA interaction was assessed by culturing the transformed yeast on SD medium lacking leucine (SD/-Leu) and supplemented with Aureobasidin A (AbA) at a minimal inhibitory concentration of 150 ng/mL for four days. Primer sequences used for cloning are provided in Supplementary Table 1.

### Histochemical localization of GUS activity

GUS activity was performed using the histochemical staining protocol described by Jefferson (Jefferson et al., 1987). Transgenic *Arabidopsis thaliana* lines with *ProCsatFT3::GUS* construct were incubated in a staining solution composed of 50 mM sodium phosphate buffer (pH 7.2), 2.0 mM potassium ferricyanide [K₃Fe(CN)₆], 2.0 mM potassium ferrocyanide [K₄Fe(CN)₆], 0.1% (v/v) and 1.0 mg/mL X-Gluc (5-bromo-4-chloro-3-indolyl-β-D-glucuronide). Samples were incubated at 37 °C for 5 hours to allow GUS expression to develop. Following staining, tissues were cleared with 70% (v/v) ethanol to remove chlorophyll and enhance contrast, then examined under a stereo microscope (ZEISS Stemi 508). To validate GUS expression patterns, at least ten independent transgenic lines were screened for the construct. Plants from three confirmed GUS-positive lines were selected for detailed expression analysis at various developmental stages, including 7, 14, and 21 days after germination (DAG), inflorescence emergence, flowering, and silique formation. For the flowering and silique stages, staining was performed on aerial tissues from soil-grown plants

### Yeast two-hybrid (Y2H) assays

Yeast two-hybrid assays were conducted following the manufacturer’s protocol using the Matchmaker™ Two-Hybrid System (Clontech). The coding sequences of CsatFD1, CsatFD2, and CsatFD3 were cloned into the bait vector pGBKT7, whereas those of CsatFT3, CsatTFL1-1, CsatTFL1-2, and CsatTFL1-3 were inserted into the prey vector pGADT7. The primers used for amplification are provided below. Bait and prey plasmids were co-transformed into the Y2HGold yeast strain and cultured on synthetic dextrose (SD) medium lacking tryptophan (SD/-Trp) and leucine (SD/-Leu) for 3–5 days at 30°C. Protein-protein interactions were assessed by selecting colonies on SD medium lacking Trp, Leu, and histidine (SD/-Trp/-Leu/-His).

### Bimolecular fluorescence complementation (BiFC)

The coding sequences of *CsatFD1*, *CsatFD2*, *CsatFD3* Cloned into YFP C-terminal (YFPC) and *CsatFT3*, *CsatTFL1-1*, *CsatTFL1-2*, *CsatTFL1-3* were cloned into the YFP N-terminal (YFPN) vectors. Primer sequences used for vector construction are provided in the supplementary table 1. These constructs were then introduced into Agrobacterium tumefaciens strain GV3101 for transient expression in 5-week-old *Nicotiana benthamiana* leaves. After co-infiltration for 48 hours, fluorescence was imaged using a confocal laser scanning microscope (Leica, Stellaris 5, Germany). YFP fluorescence was excited at 514 nm, with emission detected between 522 and 560 nm.

### Firefly Complementation Assay

The full-length coding sequences (CDSs) of CsatFT3, CsatTFL1-3, and CsatFD2 were cloned into the pCAMBIA1300-nLUC and pCAMBIA1300-cLUC vectors, respectively, and introduced into Agrobacterium tumefaciens strain GV3101. Various combinations were co-infiltrated into Nicotiana benthamiana leaves. The infiltrated plants were incubated at 22 °C for two days. Prior to LUC activity detection, the leaves were sprayed with 1 mM D-luciferin (Potassium Salt, BioVision) and incubated in the dark for 8 minutes to eliminate background fluorescence. Fluorescence was detected using a cooled CCD camera system (Bio-Rad Chemi Doc). Relative LUC intensity was quantified using ImageJ software. Statistical analysis was performed using one-way ANOVA, and significance was assessed using the Compact Letter Display (CLD) method to indicate differences at a significant p-value level. Primer sequences used in these assays are provided in Supplementary Table 1.

### Genome walker library for promoter amplification

The 5′ upstream region of the CsatFT3 gene from Crocus sativus was obtained using the Genome Walker™ kit (Clontech, USA). Genomic DNA was extracted from leaf tissue following the protocol provided with the QIAGEN DNA isolation kit, then digested with two blunt-end restriction enzymes, DraI and EcoRV, and further ligated with adapters, to generate Genome Walker libraries. Gene-specific primers (GSPs), designed based on the CsatFT3 cDNA sequence, were used alongside adaptor primers from the kit to carry out primary and nested PCR amplifications as listed in Supplementary Table 1. The amplified PCR products were resolved on a 1% (w/v) agarose gel, purified, and cloned into the pGEM-T Easy vector (Promega, USA) for sequencing. The resulting CsatFT3 promoter sequence was analyzed using the PlantCARE database (Lescot et al., 2002) to identify potential cis-regulatory elements involved in transcriptional regulation.

### Virus-induced gene silencing (VIGS)

Virus-induced gene silencing (VIGS) was carried out following the protocol established in lab (Kalia et al., 2024). Gene-specific fragments were amplified for *CsatFT3* (300 bp, flanked by EcoRI sites), *CsatFD2* (292 bp, flanked by BamHI sites), *CsatTFL1-3* (302 bp, containing both EcoRI and BamHI sites), and *CsatSVP2* (318 bp, with EcoR1and BamH1 sites) restriction sites at both termini. These fragments were cloned into the pTRV2 vector to generate the constructs. Agrobacterium-mediated infiltration was performed as previously described (Kalia et al., 2024). To confirm gene silencing, transcript levels of *CsatFT3*, *CsatFD2*, *CsatTFL1-3*, and *CsatSVP2* were quantified by RT-qPCR in apical buds collected at the floral induction stage. Phenotypic analysis of flower development was conducted when floral buds reached the floral emergence stage. Primer sequences used for the VIGS assay are listed in Supplementary table 1.

### Luciferase assay

To investigate the regulatory effect of *CsatSVP2* on the *CsatFT3* promoter, a luciferase reporter assay was conducted in *Nicotiana benthamiana* leaves. Promoter sequences of *CsatFT3* were cloned into the Gateway-compatible vector pGWB435 (Invitrogen) upstream of the luciferase (Luc) reporter gene. Three promoter constructs were generated: (i) the full-length promoter (FL), (ii) a truncated fragment containing the CArG motif(T1), and (iii) a truncated fragment lacking the CArG motif (T2). The *CsatSVP2* coding sequence was separately cloned into pGWB402 under the control of the CaMV35S promoter to serve as an effector. Control treatments included empty vectors (pGWB435 and pGWB402) as a negative control, and *ProCsatFT3*::Luc alone as a positive control. Constructs were introduced into the *Agrobacterium tumefaciens* strain GV3101 and co-infiltrated into *Nicotiana benthamiana* leaves. Each leaf was divided into four quadrants to enable side-by-side comparisons of different treatments within the same biological context. Luciferase activity was visualized and quantified using ImageJ software. The experiment was repeated independently at least three times in separate leaves to ensure reproducibility. One-way ANOVA was used for statistical analysis, and differences among treatments were determined using Compact Letter Display (CLD) to indicate statistically significant differences (p < 0.05).

### Statistical analyses

All statistical analyses were conducted using GraphPad Prism version 10 for Windows (GraphPad Software, Boston, MA, USA; https://www.graphpad.com). One-way ANOVA followed by Tukey’s Honest Significant Difference (HSD) test was applied for multiple comparisons. Statistically significant differences between groups at p < 0.05 are indicated by different letters, as determined by Compact Letter Display (CLD). Two-way ANOVA to assess the effects of temperature, month, and their interaction, followed by Sidak’s multiple comparisons test and Student’s t-test for one-to-one comparison. (*: p<0.05; **: p<0.01***: p<0.001) (****: p<0.0001).

## Results

### CsatFT3 functions as a temperature-sensitive regulator of floral induction in saffron

Our previous work identified several PEBP family genes implicated in flowering regulation in saffron, with *CsatFT3* emerging as a potential key regulator of floral induction (Kalia et al., 2023). However, that study was limited to spatial and temporal expression profiling during the reproductive phase transition. To further elucidate the functional role of *CsatFT3*, we investigated its expression dynamics across vegetative and reproductive phases in saffron apical buds. We found that *CsatFT3* is specifically expressed during the reproductive phase, with transcript accumulation coinciding with the floral induction stage (Figure 1A, Supplementary Figure 1). In contrast, its expression was barely detectable in apices of newly developing corms and during the vegetative phase in winter. Given that low temperatures are known to suppress floral induction in saffron (Jose-Santhi et al., 2023), we next assessed *CsatFT3* expression in apical buds of corms stored under low (8°C) and ambient (25°C) temperature conditions following the vegetative phase. Consistent with a temperature-responsive role, *CsatFT3* transcript levels were markedly reduced in corms stored at 8°C compared to those maintained at 25°C (Figure 1B).

**Figure 1.**
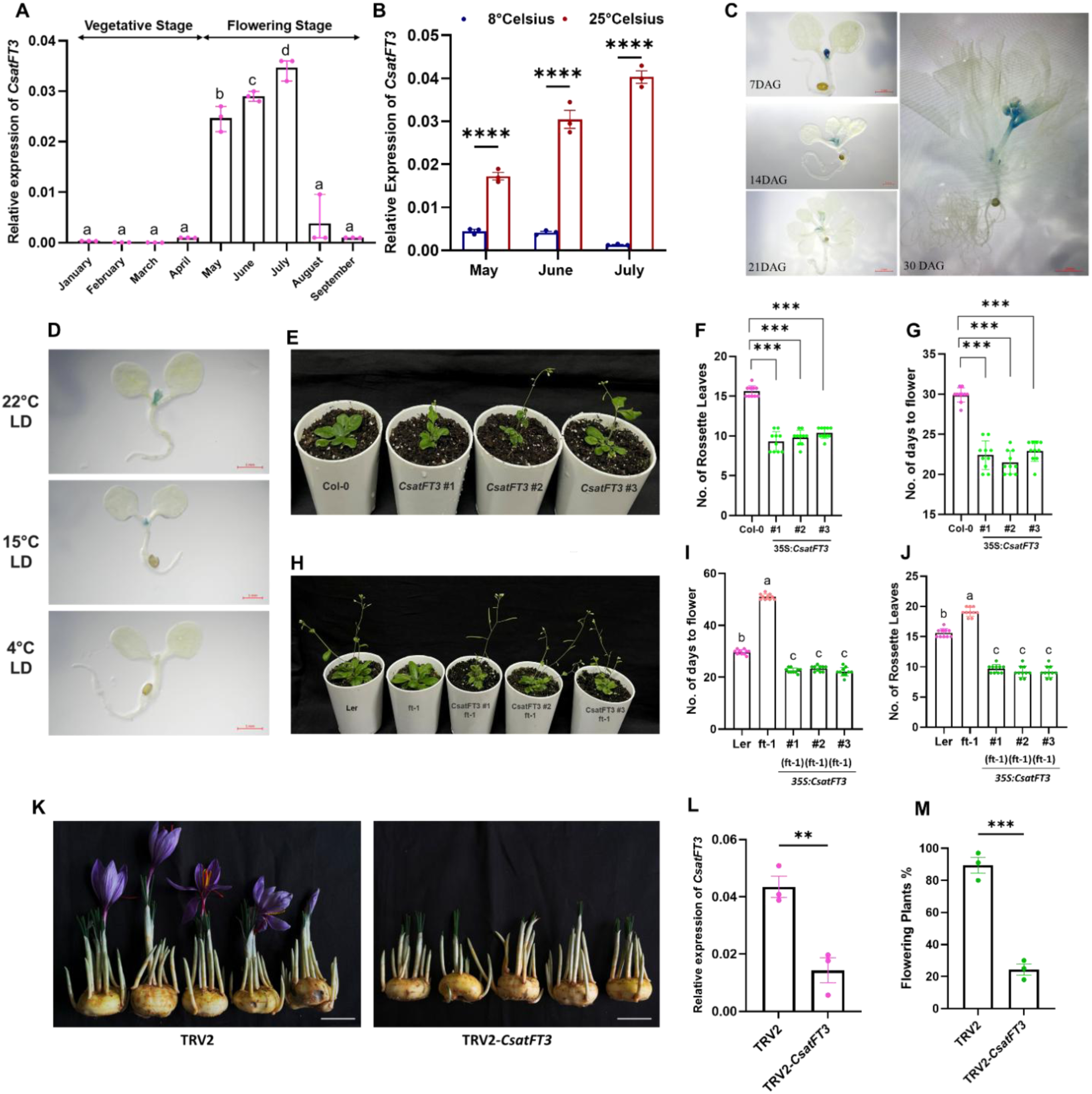
*CsatFT3* is involved in temperature mediated floral induction in saffron. **(A)** CsatFT3 transcript levels in apical buds across developmental stages show specific expression during the reproductive phase. **(B)** Relative expression of CsatFT3 in apical buds of saffron corms stored at ambient (25°C) and low (8°C) temperatures. Data are means ± SEM from 3 biological replications, where n=10 for each replicate. **(C)** Histochemical GUS assay of the *CsatFT3* promoter (proCsatFT3) in transgenic Arabidopsis in long-day conditions (LD) during various developmental stages. 1.2kb sequence upstream of the CsatFT3 translation start sites (proCsatFT3) was fused with uidA (GUS) reporter gene. Different developmental stages include Arabidopsis plants after 7 Days After Germination (DAG), 14DAG, 21DAG, and 30DAG. FT3 expression is confined to meristem and inflorescence **(D)** Temperature sensitive GUS activity at 3 different temperatures. The experiment was repeated three times independently with similar results. **(E-G)** Ectopic expression of CsatFT3 in Arabidopsis resulted in early flowering phenotype. Statistical significance was analyzed by Student’s t-test (*P < 0.05). **(H-J)** Functional complementation of the *ft-1* mutant with CsatFT3 restores flowering. Representative line expressing number of rosette leaves and number of days to flower. Total leaf number was calculated by combining total rosette leaves. Data represent a minimum of 10 plants scored for each line. Statistical significance was analyzed by Student’s t-test (*P < 0.05). **(K)** Representative flowering vs. non-flowering phenotypes in control and CsatFT3-silenced saffron corms. The scale bar represents 2 cm. **(L)** qRT-PCR validation of CsatFT3 transcript knockdown in saffron corm apices following VIGS (n=3 technical replicates, 10 independent corms). Error bars are shown as Standard Error of Mean (SEM) of three technical repeats. **(M)** Percentage of flowering in VIGS-silenced vs. TRV2 control corms. The data presented includes the mean percentage of flowering, with error bars representing the Standard Error of Mean (SEM) of three technical repeats (n=3) and for each replicate, 10 corms were used. All data in this figure are shown as mean ± SEM. Statistical Significance for Data in panel A, I, & J are analyzed using one-way ANOVA followed by Tukey’s Honest Significant Difference (HSD) test for multiple comparisons. Different letters indicate statistically significant differences between groups at p < 0.05, according to Compact Letter Display (CLD). Data in panels B are analyzed using two-way ANOVA to assess the effects of temperature, month, and their interaction, followed by Sidak’s multiple comparisons test. Asterisks indicate significant differences between 8°C and 25°C within each month (**** p < 0.0001), F, **G**, **L**, and **M** are analyzed using the student’s t-test (*: p<0.05; **: p<0.01).

To further investigate tissue-and temperature-specific expression of the *CsatFT3* gene, we isolated a 1.2 kb fragment of the *CsatFT3* promoter (Supplementary figure 2) that contains several core, distal, and proximal promoter elements and constructed a promoter-GUS fusion construct. GUS-driven transcription reporter lines of *CsatFT3* displayed meristem and flower-specific expression in Arabidopsis, consistent with our spatiotemporal expression profiling in saffron (Figure 1C and Supplementary figure 3). Furthermore, these *ProCsatFT3::GUS* fusion lines demonstrated temperature-sensitive regulation, with expression gradually decreasing as the temperature was lowered, in line with our previous findings that low temperatures suppress the expression of *CsatFT3* (Figure 1D). At low temperatures (4°C), GUS activity was nearly undetectable, indicating a significant suppression of expression under cold conditions. These findings reinforce the role of *CsatFT3* in temperature-sensitive flowering regulation in saffron and suggest that its expression is activated under the relatively high-temperature conditions that promote floral induction.

To functionally validate the role of *CsatFT3* in floral regulation, we ectopically expressed *CsatFT3* in Arabidopsis Col-0 and complemented the late-flowering *ft-1* mutant (Ler-0). Overexpression in Col-0 resulted in a significant early-flowering phenotype (Figure 1E–G), while *ft-1* plants transformed with *CsatFT3* exhibited restored or even earlier flowering compared to wild-type (Figure 1H–J). These results confirm that *CsatFT3* is functionally conserved and capable of promoting flowering in a heterologous system. Following confirmation of CsatFT3’s involvement in floral induction and flowering, finally, to evaluate the role of *CsatFT3* in its native we performed Virus-Induced Gene Silencing (VIGS) of *CsatFT3* in saffron corms and monitored flowering. Silencing of *CsatFT3* led to a significant reduction in its transcript levels in apical buds (Figure 1K) and was accompanied by a marked decrease in flowering percentage compared to TRV2 mock-infected controls (Figure 1L–M). No significant decrease of *CsatFT1* and *CsatFT2* was observed in TRV2-silenced CsatFT3 corms (Supplementary Figure 4). Together, these findings establish CsatFT3 as a temperature-sensitive floral inducer in saffron, functioning as a key regulatory component during the transition from vegetative to reproductive growth.

### Identification of florigen activation complex component FD, involved in flowering induction in saffron

*FT* like genes requires interaction with bZIP transcription factor-*FD* to form FT-FD complex that directly activates the floral meristem identity genes. Thus, to identify the FD like genes involved in floral induction of saffron we have cloned full length cDNAs for three FD-like genes (*CsatFD1*, *CsatFD2* and *CsatFD3*), which encodes for 197, 215 and 245 aa of proteins respectively. All the three FDs are similar to FD proteins from other plants and contains the conserved basic region, Leucine zipper region and the functionally important conserved threonine (T)/SAP motif at the C terminus (Supplementary Figure 5A). Phylogeny aligns CsatFD’s with monocot FDs with Asparagus officinalis as the nearest family member (Supplementary Figure 5B).

We next investigated the expression profiles of the three saffron *FD* genes during the vegetative and flowering induction stages. Interestingly, the expression of *CsatFD2* coincided with the floral induction stage, whereas *CsatFD3* showed higher expression at later stages of sprouting, specifically in July, when floral differentiation and vegetative growth begin (Figure 2 A). In contrast, *CsatFD1* was expressed at relatively stable levels throughout flower initiation and the vegetative phase, but its expression was generally lower compared to *CsatFD2* and *CsatFD3*. In saffron, the apical bud gives rise to the flower, while the axillary buds contribute only to vegetative growth, thereby distinguishing the reproductive meristem from the vegetative meristems (Kalia et al., 2022). Notably, *CsatFD2* was predominantly expressed in apical buds and flower tissues, indicating a primary role in flowering (Figure 2B, C). Additionally, *CsatFD2* showed comparatively higher expression in flower tissues compared to the other two FD genes (Figure 2C). We also examined the temperature-sensitive expression of these genes in corms stored at low (8°C) and ambient high (25°C) temperatures. Our results revealed that CsatFD2 expression was significantly downregulated at low temperatures (Figure 2D), whereas CsatFD1 and CsatFD3 did not show a similar trend (Supplementary Figure 6A). Notably, CsatFD1 expression remained unchanged at 8°C in June, the period when floral induction typically occurs. Interestingly, CsatFD3 showed higher expression in corms stored at low temperatures compared to those stored at 25°C (Supplementary Figure 6B).

**Figure 2:**
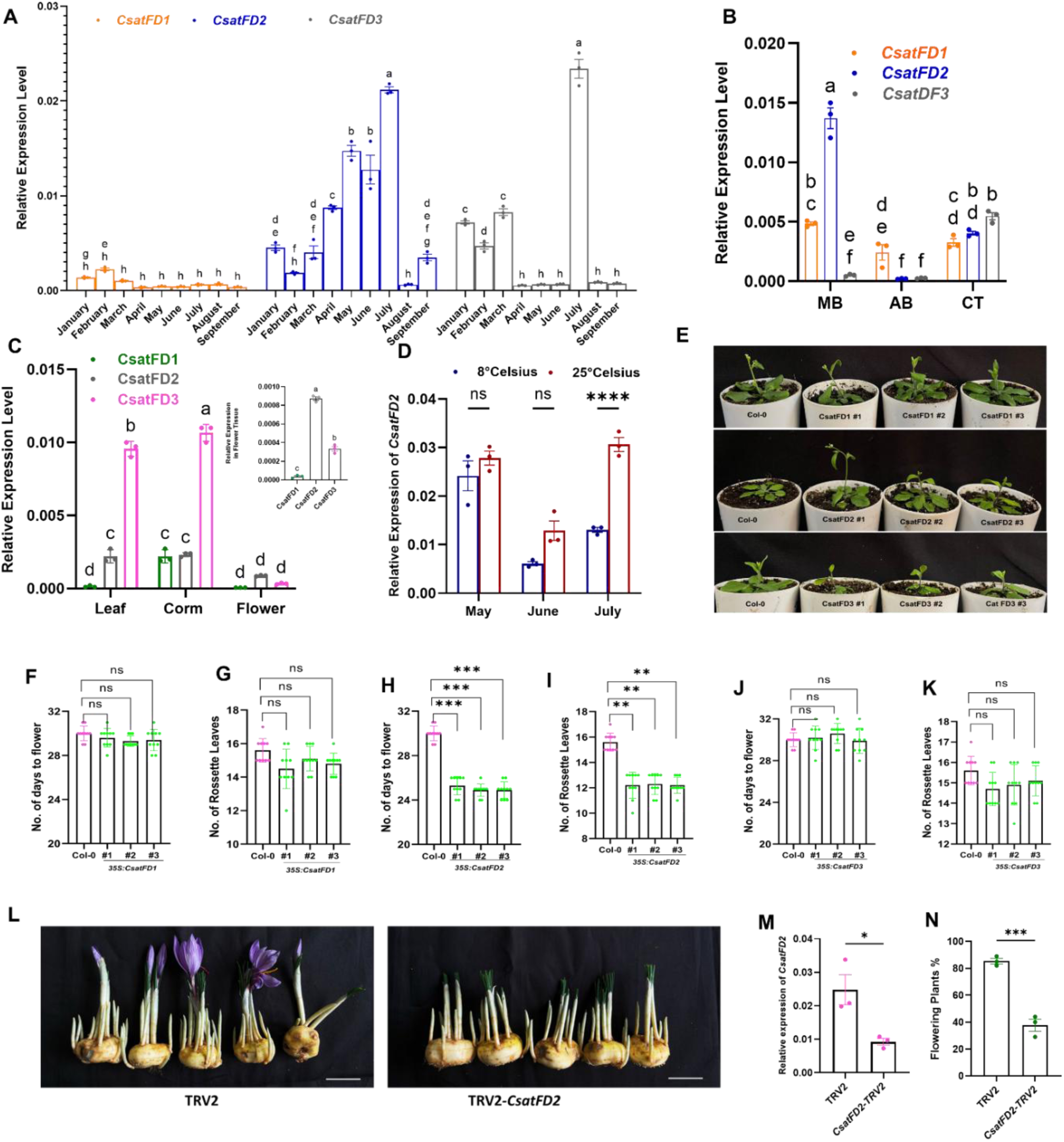
Functional characterization of FD-like genes in saffron. **(A)** Expression profiles of CsatFD1, CsatFD2, and CsatFD3 during vegetative and floral induction stages in the apical bud by RT-qPCR. Data are means ± SE of 3 independent replicates. **(B)** Expression patterns of CsatFD1, CsatFD2, and CsatFD3 in the Main bud (MB), axillary bud (AB), and corm tissue (CT) during the floral induction stage. Values are means ± SE of 3 independent replicates. **(C)** Tissue specific expression of FDs in different parts of saffron plant. **(D)** Expression analyses of *CsatFD* genes in apical buds of corms stored at 25◦ C and 8 during months of May and June (**E)** Phenotypic comparison of early flowering in CsatFD2-expressing Arabidopsis compared to control. **(F-K)** Measurements of days to flowering and rosette leaf number in CsatFD1, CsatFD2, and CsatFD3 expressing Arabidopsis plants. Values are means ± SE (n = 10). Asterisks indicate a significant difference between the transgenic Arabidopsis lines and wild-type plants (Col-0; Student’s t-test, P < 0.05). **(L)** Phenotypes of CsatFD2-TRV2 silenced lines and TRV2 lines. The scale bar represents 2 cm. **(M)** qRT-PCR validation of CsatFD2 knockdown in saffron corm apices after VIGS (n=3). Error bars are shown as SD of three technical repeats **(N)** Flowering percentage in TRV2 control and CsatFD2 VIGS-silenced saffron corms. The data presented includes the mean percentage of flowering, with error bars representing the standard error of mean (SEM) of three technical repeats (n=3) and for each replicate, 10 corms were used. All data in this figure are shown as mean ± SEM. Statistical Significance for data in panel **A** & **B** are analyzed using two-way ANOVA followed by Tukey’s Honest Significant Difference (HSD) test for multiple comparisons. Different letters indicate statistically significant differences between groups at p < 0.05, according to Compact Letter Display (CLD). Data in panel **D** are analyzed using two-way ANOVA to assess the effects of temperature, month, and their interaction, followed by Sidak’s multiple comparisons test. Asterisks indicate significant differences between 8°C and 25°C within each month (**** p < 0.0001), **F**,**G**,**H**,**I**,**J**,**K**,**M** and **N** are analyzed using the Student’s t-test (*: p<0.05; **: p<0.01).

Next, we examined the effect of ectopic expression of the saffron *FD* genes in Arabidopsis. Ectopic expression of *CsatFD2* can only induce early flowering in Arabidopsis, whereas there was no significant phenotypic change observed in *CsatFD1* and *CsatFD3* expressing Arabidopsis plants (Figure 2E). These findings were further supported by measurements of days to flowering and the number of rosette leaves in the Arabidopsis transformants (Figures 2F-K). These results further suggest that *CsatFD2* may be the primary gene involved in flowering regulation in saffron. To confirm the role of *CsatFD2* during floral induction, we silenced the gene using Virus-Induced Gene Silencing (VIGS) in flower-competent saffron corms. Compared to the TRV2 control, silencing of *CsatFD2* resulted in a reduction in flower formation and overall flowering percentage (Figure 2L). VIGS silencing of CsatFD2 also led to a significant decrease in its transcript levels (Figure 2M), but did not affect the expression of *CsatFD1* and *CsatFD3* (Supplementary Figure 7 A and B). These results suggest that *CsatFD2* is a key component of the Floral Activator Complex (FAC) involved in floral induction in saffron.

### TFL1-3/CEN1 is a newly identified negative regulator of flowering induction in saffron

In our previous study (Kalia et al., 2023), we identified two homologs of *TFL-1*/*CEN*, named Csat*TFL1-1* and Csat*TFL1-2*, and through spatio-temporal expression profiling, we proposed that TFL1-2 may play a role in determining flowering competency in saffron (Supplementary figure 8A and B). However, neither of these two *TFL-1* genes exhibited a temperature-sensitive response (Supplementary Figure 9A and B). Furthermore, ectopic expression of *CsatTFL1-1* and *CsatTFL1-2* in Arabidopsis led to delayed bolting in only *CsatTFL1-2* plants compared to wild-type (WT) under long-day (LD) conditions, whereas *CsatTFL1-1* transformants bolted normally, similar to WT plants (Supplementary figure 10A-F). Additionally, interaction studies, including BiFC and Y2H assays, did not reveal any positive interactions with *CsatFD2*, our identified positive regulator of flowering (Supplementary Figure 11A and B). These findings prompted us to investigate further, hypothesizing that another homolog of TFL1/CEN might be involved in floral induction in saffron. Through corm development studies in our lab (Jose-Santhi et al., 2023), we identified a third *TFL1* homolog, which exhibited higher expression during the vegetative phase. This gene shares similarities with *CsatTFL1-1* and *CsatTFL1-2*, but phylogenetic analysis clustered it within the CEN subgroup. Based on this, we named it *CsatTFL1-3*/*CEN1* (Supplementary Figure 12A and B). Like other TFL1/CEN family members, TFL1-3/CEN1 contains the conserved domains characteristic of this gene family. Quantitative PCR (qPCR) expression profiling revealed that *CsatTFL1-3*/*CEN1* expression is elevated during the vegetative phase, with a significant decrease during the reproductive phase (Figure 3A). Tissue-specific expression analysis confirmed that *CsatTFL1-3*/*CEN1* is predominantly expressed in the axillary bud tissue, further supporting its potential role in negatively regulating flowering (Figure 3 C and D).

**Figure 3:**
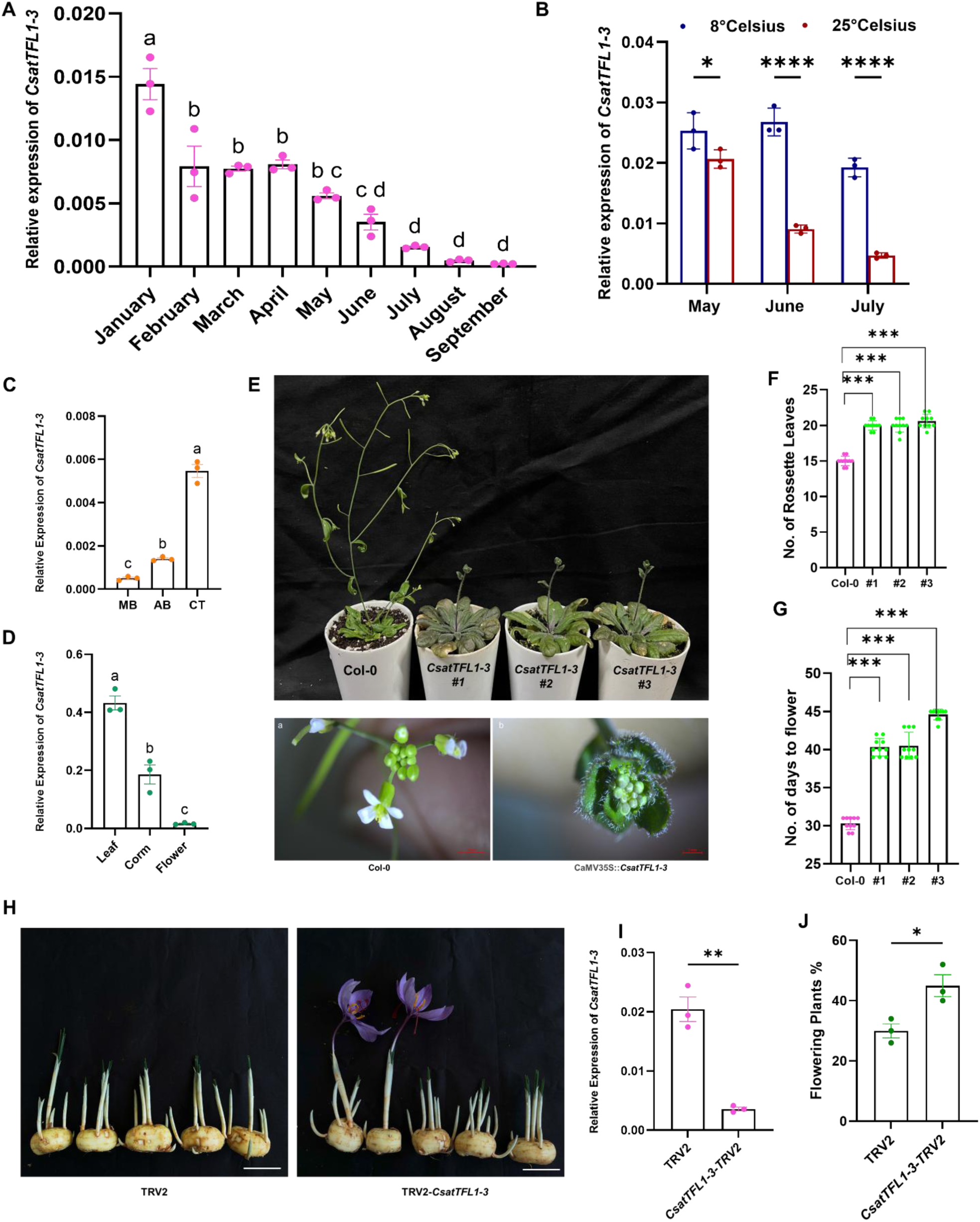
CsatTFL1-3 is negative regulator of floral induction in saffron. **(A)** Relative expression of CsatTFL1-3 during vegetative and reproductive phase in the apical bud by RT-qPCR. Data are means ± SE of 3 independent replicates. **(B)** Expression analyses of CsatTFL1-3 gene in apical buds of corms stored at 25°C and 8 °C during floral induction time **(C)** Corm tissue specific expression of TFL1-3 during floral induction. **(D)** Tissue Specific Expression of CsatTFL1-3. **(E)**Phenotypic analysis of Arabidopsis Col-0 plants ectopically expressing TFL1-3/CEN1, Hyper vegetive phase phenotype observed in CsatTFL1-3 transgenic lines of Arabidopsis. **(F and G)** Rosette leaves number and number of days to flower of 35S::TFL1-3 transgenic and control plants respectively. Values are means ± SEM (n = 10). Asterisks indicate a significant difference between the transgenic Arabidopsis lines and wild-type plants (Col-0; Student’s t-test, P < 0.05). **(H)** Virus-Induced Gene Silencing (VIGS) of TFL1-3/CEN1 in low-flower-competent saffron corms. The scale bar represents 2 cm. **(I)** The expression of *CsatTFL1-3* in *CsatTFL1-3*-TRV2 silenced lines and TRV2 lines (n=3). **(J)** Comparison of flowering percentages in VIGS-silenced versus control saffron corms. The data presented includes the mean percentage of flowering of three technical repeats (n=3) and for each replicate, 10 corms were used. All data in this figure are shown as mean ± SEM. Statistical Significance for Data in panel **A**, **C** & **D** are analyzed using one-way ANOVA followed by Tukey’s Honest Significant Difference (HSD) test for multiple comparisons. Different letters indicate statistically significant differences between groups at p < 0.05, according to Compact Letter Display (CLD). Data in panel B are analyzed using two-way ANOVA to assess the effects of temperature, month, and their interaction, followed by Sidak’s multiple comparisons test. Asterisks indicate significant differences between 8°C and 25°C within each month (**** p < 0.0001), F,G,I, & J are analyzed using the Student’s t-test (*: p<0.05; **: p<0.01).

Next, we examined the expression of *CsatTFL1-3*/*CEN1* in corms stored at low (8°C) and ambient (25°C) temperatures. Interestingly, there was no reduction in *TFL1-3*/*CEN1* expression in corms stored at low temperatures compared to those stored at ambient high temperatures, suggesting that it may be regulated by temperature and could play a role in temperature-mediated floral induction in saffron (Figure 3B). We then performed ectopic expression of *CsatTFL1-3*/*CEN1* in *Arabidopsis thaliana* (Col-0) under the control of the constitutive 35S promoter. Transgenic lines exhibited a significant delay in flowering compared to wild-type plants, as evidenced by increased days to bolting and a higher number of rosette leaves at flowering (Figure 3F-G). These results confirm the ability of TFL1-3/CEN1 to act as a floral repressor in Arabidopsis, consistent with its proposed function in saffron. Notably, ectopic expression of TFL1-3/CEN1 also led to a hyper-vegetative shoot phenotype, characterized by the development of an inflorescence composed of leaf primordia surrounded by small, serrated leaves—a phenotype not observed in TFL1-1 or TFL1-2 expressing Arabidopsis lines (Figure 3E). Finally, we performed Virus-Induced Gene Silencing (VIGS) of *CsatTFL1-3*/*CEN1* in low-flower-competent saffron corms. In comparison to TRV2 control plants, silencing of *CsatTFL1-3*/*CEN1* led to the formation of floral structures in 7-9 g corms (classified as low-flower-competent), and these flowers appeared earlier than in TRV2 controls (Figure 3G-I). Importantly, VIGS of *CsatTFL1-3*/*CEN1* did not affect the transcript levels of the other two *TFL1* homologs, TFL1-1 and TFL1-2 (Supplementary Figure 13 A and B), suggesting TFL1-3/CEN1 and not TFL1-1 or TFL1-2 could play a role in flowering in saffron. This observation suggests that TFL1-3/CEN1 acts as a negative regulator of floral induction, and its silencing promotes floral transition in otherwise low-flowering competence corms.

### The FT-FD-TFL complex involved in floral regulation

Our earlier analyses identified *CsatFT3*, *CsatFD2*, and *CsatTFL1-3*/*CEN1* as potential core components of a Floral Activator Complex (FAC) in saffron. To investigate whether these proteins physically interact to form functional regulatory complexes, we employed two complementary approaches: yeast two-hybrid (Y2H) assays and bimolecular fluorescence complementation (BiFC) in *Nicotiana benthamiana*. In the Y2H assays, the coding sequences (CDS) of the three *CsatFD* genes were cloned into the pGADT7 prey vector, and co-transformed into yeast with either *CsatFT3* or *CsatTFL1-3*/*CEN1* fused to the pGBKT7 bait vector. Interaction screening revealed that both CsatFT3 and TFL1-3/CEN1 specifically interacted with CsatFD2, but not with CsatFD1 or CsatFD3 (Figure 4 A and B). These results suggest that CsatFD2 may serve as a central interacting hub for both activator (FT3) and repressor (TFL1-3) proteins within the FAC.

**Figure 4.**
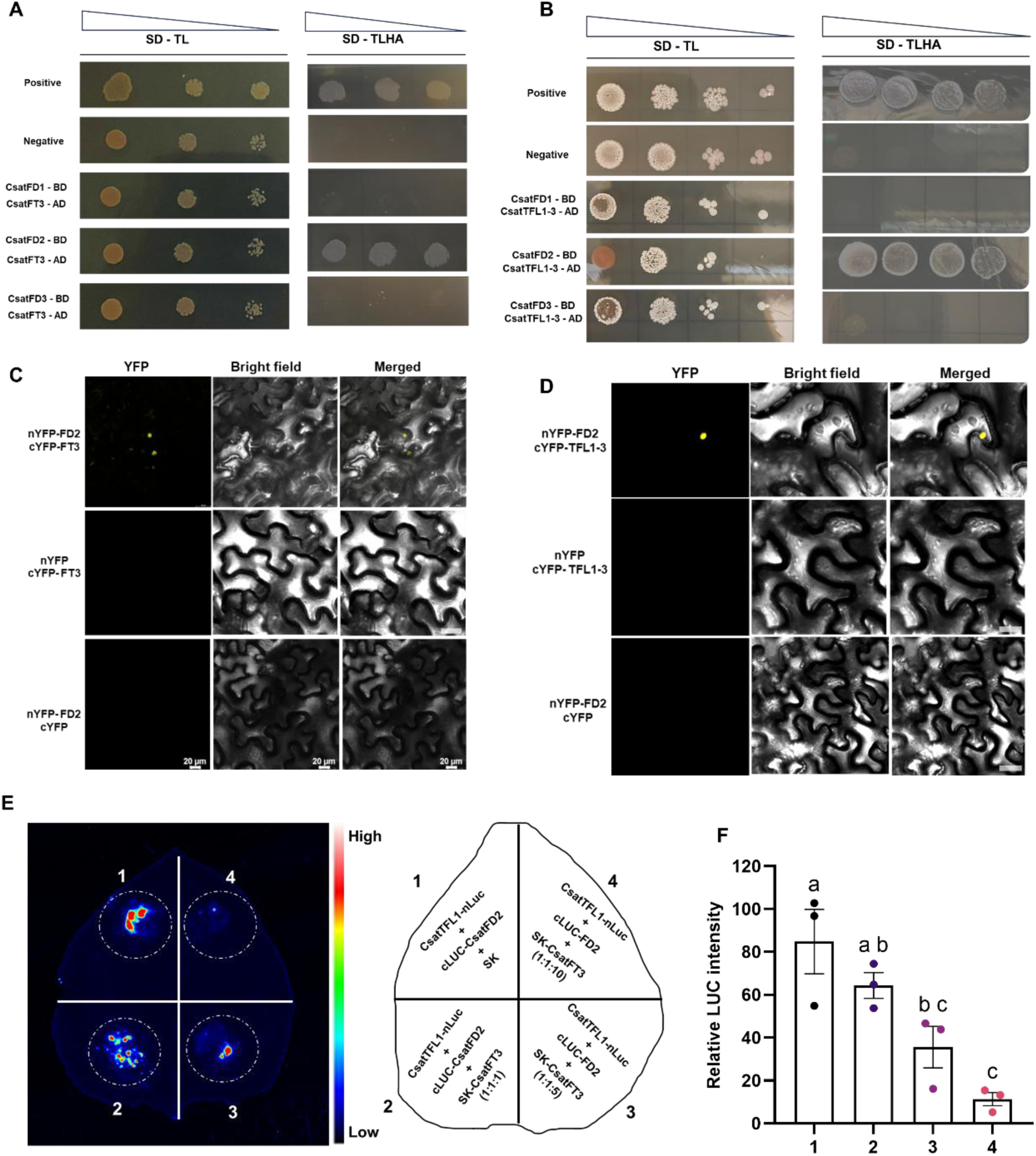
Interaction analysis of CsatFT3, CsatTFL1-3, and CsatFD2. **(A, B)** Yeast two-hybrid (Y2H) assays showing physical interactions between CsatFT3/CsatTFL1-3 and CsatFD2. The coding sequences of *CsatFT3* and *CsatTFL1-3* were fused to the activation domain in the pGADT7 vector, and *CsatFD2* was fused to the DNA-binding domain in the pGBKT7 vector. Yeast transformants were grown on double dropout (DDO) medium lacking tryptophan and leucine (selection for transformation) and quadruple dropout (QDO) medium lacking tryptophan, leucine, histidine, and adenine (selection for interaction). pGBKT7-53 and pGADT7-T were used as positive control; pGBKT7-lam and pGADT7-T interactions served as the negative control. **(C-D)** Bimolecular fluorescence complementation (BiFC) assay showing nuclear localization of the interactions between CsatFT3 or CsatTFL1-3 with CsatFD2 in *Nicotiana benthamiana* leaf epidermal cells. CsatFD2 was fused to the C-terminal half of YFP (cYFP), and CsatFT3 or CsatTFL1-3 were fused to the N-terminal half of YFP (nYFP). YFP signal indicates interaction. Bars = 20 μm. **(E)** Competitive luciferase complementation assay demonstrating the competition between CsatFT3 and CsatTFL1-3 for binding to CsatFD2 in *N. benthamiana* leaves. CsatTFL1 was fused to the N-terminal half of luciferase (nLUC), and CsatFD2 to the C-terminal half (cLUC). Increasing amounts of CsatFT3 (non-tagged, driven by 35S promoter, in SK vector) were co-infiltrated at different ratios. Images represents three independent replications. **(F)** Relative luminescence intensities indicate competition for FD2 binding with TFL1-3 and FT3. Data represented is mean of three replicates. Statistical Significance are analyzed using one-way ANOVA followed by Tukey’s Honest Significant Difference (HSD) test for multiple comparisons. Different letters indicate statistically significant differences between groups at p < 0.05, according to Compact Letter Display (CLD).

We further validated these interactions by BiFC assays in *Nicotiana benthamiana* leaves. YFP fluorescence was observed in nuclei of cells co-expressing the C-terminal half of YFP fused to CsatFD2 (cYFP–CsatFD2) with the N-terminal half of YFP fused to either CsatFT3 (nYFP–CsatFT3) or CsatTFL1-3 (nYFP–CsatTFL1-3) (Figure 4 C and D). No YFP fluorescence was detected in negative controls, including co-expression of cYFP–CsatFD2 with nYFP alone or cYFP alone with nYFP–CsatFT3 or CsatTFL1-3. Furthermore, consistent with the Y2H results, no BiFC signal was observed when CsatFT3 or TFL1-3/CEN1 were co-expressed with CsatFD1 or CsatFD3 (Supplementary Figure). Together, these findings strongly support a model in which CsatFD2 specifically interacts with both CsatFT3 and TFL1-3/CEN1, suggesting that competitive complex formation may modulate floral transition in saffron.

Given the antagonistic roles of CsatFT3 and CsatTFL1-3 in the regulation of flowering, and their respective interactions with the transcription factor CsatFD2, it was hypothesized that CsatFT3 and CsatTFL1-3 may compete for binding to CsatFD2. To investigate this, a luciferase complementation imaging (LCI) assay was conducted in *Nicotiana benthamiana* leaves to evaluate the interaction strength between CsatFD2-nLUC and cLUC-CsatTFL1-3 in the presence or absence of SK-CsatFT3.Strong luciferase (LUC) activity was observed when CsatFD2-nLUC and cLUC-CsatTFL1-3 were coexpressed, indicating a robust interaction. However, coexpression with SK-CsatFT3 significantly reduced LUC activity, suggesting that CsatFT3 interferes with the interaction between CsatTFL1-3 and CsatFD2. Moreover, this suppression was dose-dependent, with increasing amounts of CsatFT3 leading to a progressive decrease in LUC signal (Figure 4E). Biological replicates supporting these observations are presented in Supplementary figure 15 A and B. These results strongly suggest that CsatFT3 competes with CsatTFL1-3 for interaction with CsatFD2, which interferes with CsatTFL1-3’s ability to interact with CsatFD2.

### CsatSVP2 Acts Upstream of FT3 in Temperature-Mediated Floral Repression

To identify upstream genes potentially involved in temperature-dependent floral induction in saffron, we screened transcriptome data representing the suppression of floral induction under low temperatures (Jose-Santhi et al., 2023) and identified two partial SVP-like MADS-box genes that were upregulated in non-flowering samples. SVP-like genes are known to regulate flowering in a temperature-dependent manner (Lee et al., 2007). Using long-read and other in-house transcriptome data, and based on sequence similarity, we obtained full-length sequences of these two SVP-like genes. We isolated and cloned them from saffron, naming them *CsatSVP1* and *CsatSVP2*. Sequence analysis indicated that while these genes are highly similar, they code for different amino acid sequences. Multiple sequence alignment revealed that both CsatSVP1 and CsatSVP2 contain the conserved MADS-box domain, and phylogenetic analysis grouped them with SVP-like genes from other geophytes, such as Narcissus, Zingiber, and Lilium (Supplementary Figure 16 A and B).

To further characterize their role in flowering, we performed expression analysis of *CsatSVP1* and *CsatSVP2* during vegetative and floral induction stages. Both genes showed a significant reduction in expression during the floral induction stage (June); however, the reduction in *CsatSVP2* was more pronounced and correlated closely with floral induction and subsequent stages (Figure 5A). We then evaluated their expression in corms stored under floral inductive (25°C) and non-inductive (8°C) conditions. *CsatSVP2* transcript levels remained high at both temperatures, suggesting it may contribute to the floral repression observed at low temperatures (Figure 5C). In contrast, *CsatSVP1* expression decreased under both conditions, showing no temperature-dependent pattern (Figure 5B). Additionally, *CsatSVP2* expression was comparatively higher in floral tissues than *CsatSVP1*, further supporting its potential regulatory role (Supplementary Figure 17).

**Figure 5:**
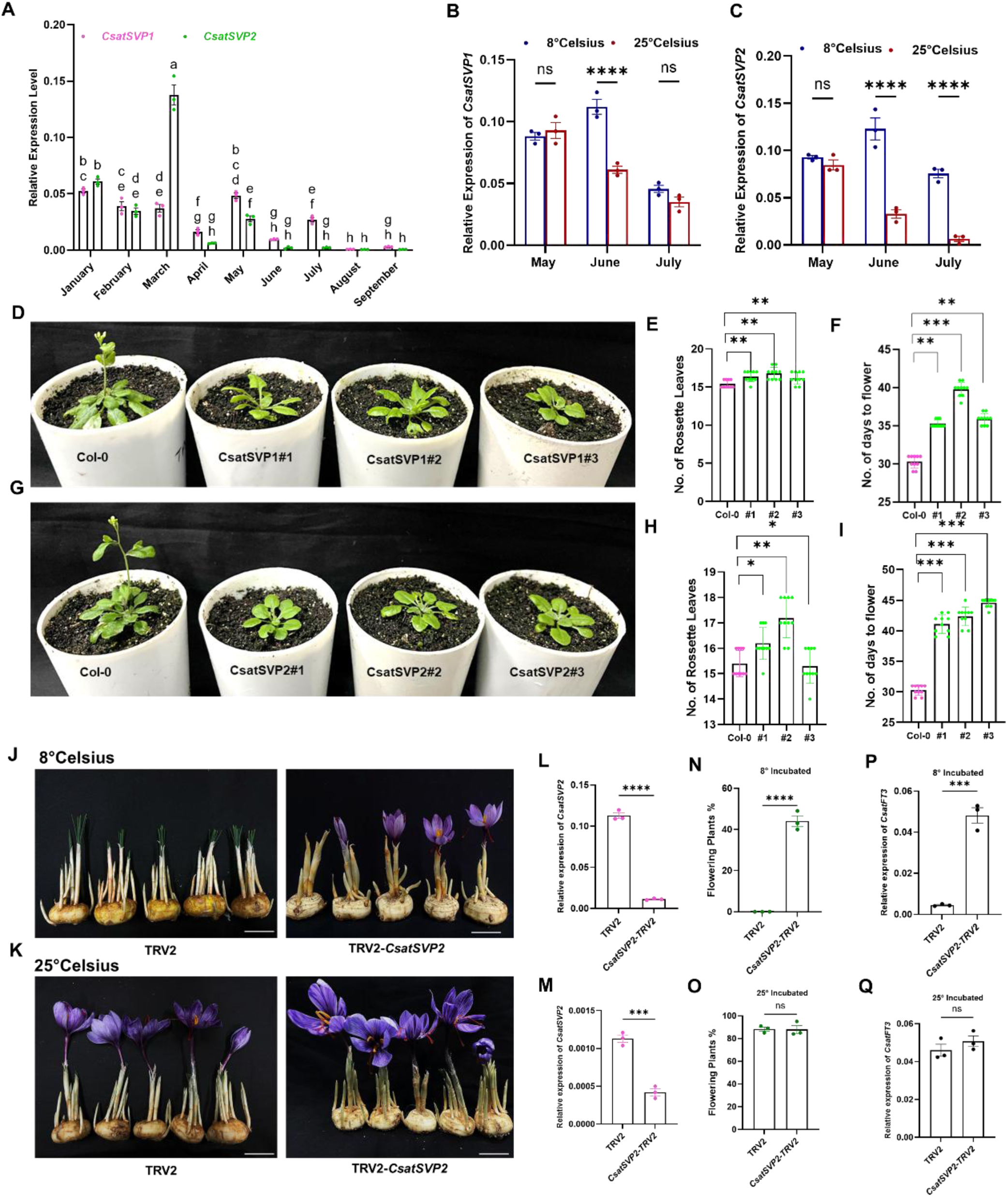
Functional characterization of CsatSVP1 and CsatSVP2 in temperature-dependent floral regulation. **(A)** Expression profiling of CsatSVP1 and CsatSVP2 in apical bud during different vegetative and flowering stages. Data are means ± SE from 3 replications. **(B and C)** Expression of SVP1 and SVP2 genes in corms stored at inductive (25°C) and non-inductive (8°C) temperatures respectively. **(D)** Flowering phenotypes of Arabidopsis lines ectopically expressing CsatSVP1. **(E and F)** Rosette leaves number and number of days to flower of CsatSVP1 transgenic and control plants respectively **(G)** Phenotypic changes of the floral organs in 35S::CsatSVP2 transgenic Arabidopsis plants. **(H-I)** Rosette leaves number and number of days to flower of CsatSVP2 transgenic and control plants respectively. The total leaf number was calculated by combining total rosette leaves. **(J-K)** Silencing of CsatSVP2 by VIGS rescues flowering under low-temperature conditions in saffron. The scale bar represents 2 cm. **(L-M)** qRT-PCR validation of CsatSVP2 knockdown in saffron corm apices after VIGS stored at different temperatures (n=3). Error bars are shown as SD of three technical repeats. **(N-O)** Flowering percentage in TRV2 control and CsatSVP2 VIGS-silenced saffron corms stored at low (8°C) and ambient (23°C) temperatures. The data presented includes the mean percentage of flowering, with error bars representing the standard deviation (SD) of three technical repeats (n=3) and for each replicate, 10 corms were used. **(P-Q)** Relative expression of CsatFT3 in SVP2-silenced versus control corms under different temperature regimes. All data in this figure are shown as mean ± SEM. Statistical Significance for data in panel A are analyzed using two-way ANOVA followed by Tukey’s Honest Significant Difference (HSD) test for multiple comparisons. Different letters indicate statistically significant differences between groups at p < 0.05, according to Compact Letter Display (CLD). Data in panel B & C are analyzed using two-way ANOVA to assess the effects of temperature, month, and their interaction, followed by Sidak’s multiple comparisons test. Asterisks indicate significant differences between 8°C and 25°C within each month (**** p < 0.0001), E, F, H, I, L, M, N, O, P & Q are analyzed using the student’s t-test (*: p<0.05; **: p<0.01).

Next, to functionally validate these genes, we ectopically expressed both *CsatSVP1* and *CsatSVP2* in Arabidopsis under a constitutive promoter. Ectopic expression of both genes resulted in delayed flowering, with a more pronounced delay in the CsatSVP2-transformed lines compared to both the control and *CsatSVP1* lines (Figure 5D-I). Arabidopsis plants expressing *CsatSVP2* displayed various phenotypic changes in flower bud and flower morphology, producing smaller, more compact flowers and shorter siliques (Supplementary Figure 18). To investigate the role of *SVP* in temperature-mediated floral suppression, we silenced *CsatSVP2* in saffron corms using Virus-Induced Gene Silencing (VIGS). Csat*SVP2* was selected due to its expression pattern and phenotype in Arabidopsis, which correlated with its expected function. Since low temperatures during storage or dormancy are known to suppress flowering, and CsatSVP2 may be involved in this process, we stored the *CsatSVP2*-silenced corms at low temperatures, alongside a mock control (TRV2). Silencing of SVP2 resulted in flowering in corms at low temperatures, whereas the TRV2-treated corms stored at low temperatures did not flower (Figure 5J). In parallel, corms stored at 25°C—an inductive condition—showed flowering in both control and SVP2-silenced groups, confirming that *CsatSVP2* acts specifically in the floral suppression mechanism under cold conditions (Figure 5K). Gene expression analysis further revealed that *CsatSVP2* silencing prevented the typical low-temperature-mediated suppression of *CsatFT3* expression, thereby facilitating flowering (Figure 5P and Q). These findings suggest that CsatSVP2 and CsatFT3 act within the same regulatory pathway, mediating temperature-responsive floral induction in saffron.

### SVP2 Directly Binds to the CsatFT3 Promoter and Represses Its Expression

Our previous results indicated that CsatSVP2 and CsatFT3 function in the same temperature-sensitive regulatory pathway: low temperature maintains or upregulates CsatSVP2 expression, which in turn suppresses *CsatFT3*, thereby repressing floral induction. Notably, *CsatFT3* transcript levels were restored in *CsatSVP2*-silenced corms, suggesting that CsatSVP2 functions upstream of *CsatFT3*. To further explore the mechanism of this regulation, we analyzed the CsatFT3 promoter sequence to identify potential binding motifs for MADS-box transcription factors. In silico analysis of the 1.2 kb FT3 promoter, previously shown to drive temperature-responsive expression, revealed a conserved CArG-box motif (CCATTTAAGG) located approximately 827 bp upstream of the ATG start codon that is established for binding of MADS boxes genes including SVP orthologs (Figure 6A). To experimentally test further whether CsatSVP2 directly binds to the *CsatFT3* promoter, we performed a yeast one-hybrid (Y1H) assay. The Y1H assay confirmed that CsatSVP2 specifically binds to the *CsatFT3* promoter, supporting the hypothesis of direct regulation (Figure 6B). To further validate the functional specificity of this interaction, we employed a luciferase reporter assay using *Nicotiana benthamiana* transient expression (Figure 6C) and conducted a luciferase reporter assay using three different FT3 promoter fragments: (1) the full-length promoter (∼1.2kb), (2) a truncated version containing the conserved CArG motif (T1), and (3) a truncated version lacking the motif (T2) (Figure 6D). Co-expression of CsatSVP2 with these constructs in *Nicotiana benthamiana* revealed that luciferase activity was significantly reduced in both the full-length and T1 constructs (Figure6 D and F). In contrast, no repression was observed with the T2 construct, confirming that SVP2 specifically suppresses FT3 expression through direct binding to the CArG motif (Figure 6F). Together, these results establish a mechanistic model in which CsatSVP2 acts as a temperature-responsive floral repressor that directly binds to the FT3 promoter via a conserved CArG motif, thereby repressing FT3 expression and floral induction under low-temperature conditions.

**Figure 6.**
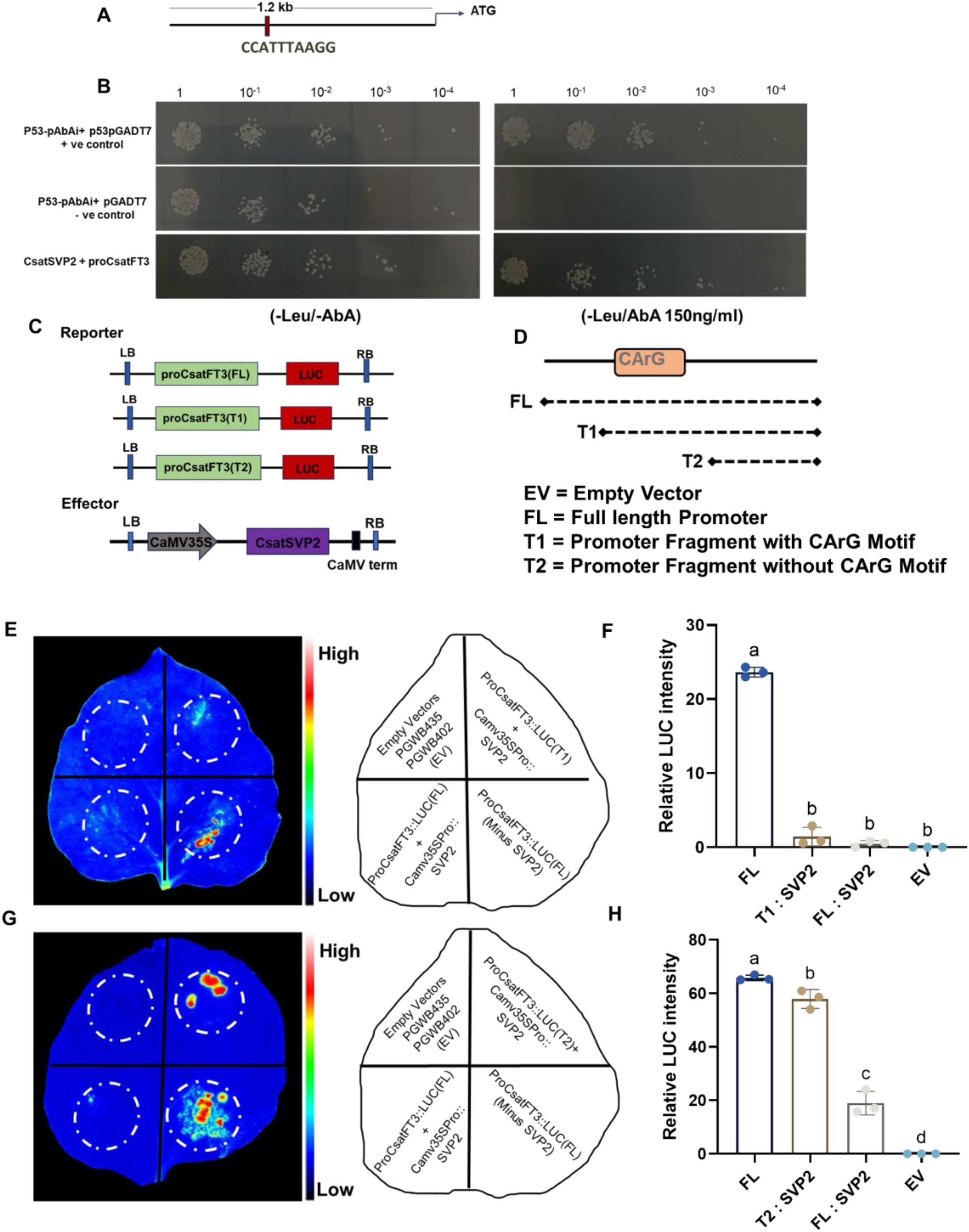
CsatSVP2 directly binds to the FT3 promoter via the CArG motif and represses its transcription. **(A)** Schematic diagram of the 1.2 kb CsatFT3 promoter showing the location of a conserved CArG-box motif (CCATTTAAGG) approximately 827 bp upstream of the ATG start codon. **(B)** Yeast one-hybrid assay showing interaction between CsatSVP2 and the FT3 promoter region containing the CArG motif. Growth on selective media confirms binding specificity. **(C)** Schematic diagram of the effectors and reporters used in dual-luciferase reporter. **(D)** Schematic of the dual-luciferase constructs used for transient expression in *Nicotiana benthamiana*: FL (Full length promoter) T1 (with CArG motif) and T2 (without). **(E)** Relative luciferase activity in leaves co-expressing CsatSVP2 with the full promoter and T1 fragment construct of *CsatFT3* gene. **(F)** Relative luciferase activity in leaves co-expressing CsatSVP2 and CsatFT3 full promoter and with T2 fragment constructs. Significant repression is observed only in the T1 construct, indicating that CsatSVP2 represses *CsarFT3* transcription through the CArG motif. **(G and H)** Quantification of luciferase activity. Bars represent mean ± SEM (n = 3). All data in figure are shown as mean ± SEM. Statistical Significance for Data in panel **F** & **H** areanalyzed using one-way ANOVA followed by Tukey’s Honest Significant Difference (HSD) test for multiple comparisons. Different letters indicate statistically significant differences between groups at p < 0.05, according to Compact Letter Display (CLD)

## Discussion

Saffron is a high-value crop cultivated in limited regions due to its strict temperature requirements, which distinctly regulate its vegetative and reproductive growth. High temperatures induce flowering, while subsequent cooling triggers flower formation. In contrast, low temperatures suppress floral induction and support vegetative growth, enabling overwintering and daughter corm development (Molina et al., 2005a). This temperature sensitivity restricts saffron cultivation to specific climates. Although environmental control of flowering has enabled cultivation in soilless and controlled settings, the molecular mechanisms underlying temperature-regulated flowering remain poorly understood. As a sterile crop, saffron cannot be improved through conventional breeding, making biotechnological interventions essential. However, limited genomic and genetic resources, challenges in genetic transformation, and a scarcity of molecular studies have hindered progress in functional genomics, creating a critical knowledge gap that must be addressed.

In this study, we have identified a temperature-sensitive transcriptional network that regulates flowering in saffron, uncovering both conserved and saffron-specific components involved in floral induction (Figure 7). Our data establish a regulatory module involving CsatFT3, CsatFD2, CsatTFL1-3/CEN1, and CsatSVP2 as key components in mediating floral transition in response to temperature cues. Although the core elements of the Florigen Activation Complex (FAC) - FT, FD, and TFL1 are conserved across angiosperms, their expression patterns, protein interactions, and functional dynamics in saffron suggest neofunctionalization in line with its geophytic lifecycle. For instance, Notably, CsatFT3 and CsatFD2 are upregulated at higher temperatures and co-expressed in the floral meristem, promoting flowering. In contrast, other FT and FD homologs show no such expression correlation (Figure 2; (Kalia et al., 2022), indicating their limited or no involvement in floral induction. Similarly, CsatTFL1-3 functions as a floral repressor by competitively binding to CsatFD2, whereas the other two TFL1 homologs neither interact with CsatFD2 nor delay flowering in Arabidopsis, suggesting a lack of functional redundancy and functional specificity. Functional validation in both Arabidopsis and saffron confirms that floral induction is governed by the dynamic interplay between CsatFT3, CsatFD2, and CsatTFL1-3.Given the roles of gene duplication, neofunctionalization, and lineage-specific gene expansion in driving the specialization of reproductive and vegetative pathways in geophytes (Khosa et al., 2021), our identification of distinct regulatory factors and their molecular interactors from multigene families in saffron represents a significant advance.

**Figure 7.**
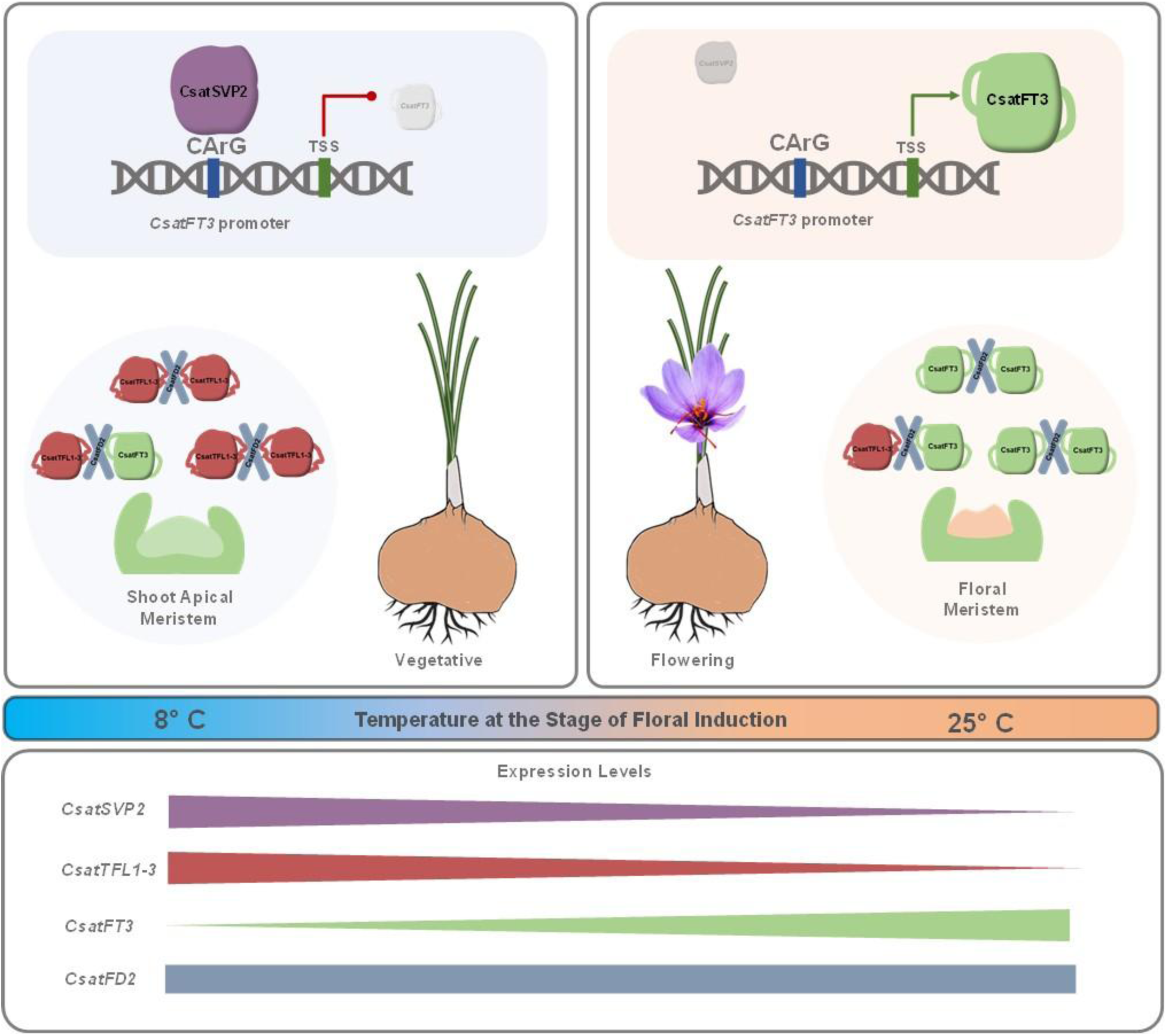
Proposed model of temperature-mediated flowering regulation in saffron. This schematic model illustrates the temperature-regulated floral induction pathway in saffron, emphasizing the key roles of CsatFT3, CsatFD2, TFL1-3/CEN1, and CsatSVP2 in transitioning from vegetative to reproductive growth. At low temperatures (e.g., below 10°C), *CsatFT3* expression remains low, and TFL1-3/CEN1 binds to CsatFD2, preventing the formation of the Floral Activator Complex (FAC) and maintaining the shoot apical meristem in a vegetative state and inhibiting flowering. Additionally, CsatSVP2 is upregulated at low temperatures and binds directly to a conserved CArG motif in the CsatFT3 promoter, repressing its transcription and reinforcing the suppression of flowering. As temperatures increase beyond a critical threshold (e.g., above 25 °C), CsatFT3 expression is upregulated, enabling its interaction with CsatFD2 to form the FAC, which in turn activates floral meristem identity genes and promotes floral initiation. Concurrently, TFL1-3/CEN1 expression is downregulated, releasing the repression on CsatFD2 and facilitating the transition to reproductive development.

A distinguishing feature of floral induction observed in saffron is the spatial restriction of flowering-related gene expression predominantly to the meristem. All the genes identified in this study were specifically expressed in the apical (flowering) meristem. During floral induction, saffron corms are either underground or stored in the dark and remain leafless, indicating that the molecular signals initiating flowering are localized within the apical buds. Such meristem-localized control may reflect an adaptation to the geophytic habit of saffron, which undergoes floral induction in the absence of foliage, similar to what has been observed in other geophytes such as tulips (Leeggangers et al., 2017a). This localization supports the hypothesis that temperature cues are sensed and interpreted locally within the corm, rather than systemically via leaves, as observed in many other species, including some geophytes (Lee et al., 2013b; Navarro et al., 2011; Wigge, 2011; Wu et al., 2022; Yan et al., 2021).

Temperature plays a pivotal role in regulating flowering time across plant species, often through conserved thermosensory pathways. In Arabidopsis, warmer temperatures promote flowering via *FT* activation (Balasubramanian et al., 2006). while in crops like strawberry and onion, cold exposure induces flowering by triggering FT-like genes (Heide, 1977; Lee et al., 2013b). Interestingly, the effect of temperature on flowering is not always straightforward. Temperatures above 25°C seem to have both inductive and repressive effects on flowering, depending on the species and environmental conditions (Nakano et al., 2013; Noy-Porat et al., 2013; Oda et al., 2012). This temperature sensitivity is closely linked to the regulation of FT, a key gene that integrates various environmental signals and controls the timing of flowering. In *Narcissus tazetta* (daffodils), floral induction is triggered by high ambient temperatures (25°C) through the activation of *NtFT*, while in tulips, *TgTFL1* downregulation and *TgFT2* upregulation contribute to temperature-induced floral induction (Leeggangers et al., 2017b; Noy-Porat et al., 2013). In contrast, in species like onion and lily, which flower in spring, vernalization (low-temperature treatment) is required to activate the expression of *AcFT2* and *LiFT*, respectively (Lee et al., 2013b). In our study, we also found that *CsatFT3*, a key flowering-time gene in saffron, is temperature-regulated, with its expression being induced by high ambient temperatures and suppressed by low temperatures (Figure 1B). Similarly, *CsatFD2*, a partner of *CsatFT3* in the flowering regulatory network, follows a similar expression pattern, suggesting their involvement in the temperature-mediated floral induction pathway in saffron. Interestingly, we observed that *CsatTFL1-3*, a gene that maintains the vegetative state and prevents early flowering, exhibits a reciprocal expression pattern. In Arabidopsis low and ambient temperature regulation of flowering involves the gene *TFL1* (Strasser et al., 2009). Higher expression of *CsatTFL1-3* is observed during the vegetative phase, which coincides with the low winter temperatures during saffron’s growth cycle (Figure 3A). This expression pattern is crucial for maintaining the vegetative phase and delaying flowering, as confirmed by the hyper-vegetative shoot phenotype in *TFL1*-3 overexpressing Arabidopsis plants (Figure 3) similar to what was observed when Arabidopsis *TFL1* is overexpressed (Lee et al., 2019). In contrast, as temperatures rise in the summer, the expression of CsatFT3 increases (Figure 1A), likely triggering the activation of floral pathway genes and the transition to reproductive development. Overall, a balance between CsatFT3, CsatTFL1-3, and their interaction with CsatFD2 governs floral induction in a temperature-dependent manner. This delicate interplay between temperature and gene expression is vital for ensuring proper timing of flowering in saffron, aligning the plant’s reproductive phase with favourable environmental conditions.

We also identify CsatSVP2, a MADS-box transcription factor, as a critical floral repressor acting upstream of FT–FD–TFL core module. MADS-box genes are key regulators of flowering time, with SVP-like genes playing central roles in ambient temperature-mediated floral control (Jin and Ahn, 2021). CsatSVP2 directly binds a conserved CArG-box in the CsatFT3 promoter, repressing its expression under low temperatures. Silencing of SVP2 alleviated low-temperature-induced flowering suppression and restored *CsatFT3* expression, thereby establishing a direct mechanistic link between ambient temperature perception and floral induction (Figure 5). In *Arabidopsis thaliana*, a comparable regulatory pathway has been described, where SVP negatively regulates FT expression by directly binding to a CArG motif in its promoter (Lee et al., 2007). Additionally, SVP levels are known to increase under low temperatures and are degraded at higher temperatures, facilitating early flowering as SVP abundance declines (Blázquez et al., 2002; Lee et al., 2013a; Sureshkumar et al., 2016). Interestingly, individual members of the SVP gene family have been shown to play distinct roles in processes such as bud dormancy, flowering initiation, and floral development reflecting adaptations of developmental timing to local environmental conditions and seasonal temperature fluctuations. In tulips, gene expression analyses have similarly identified *TgMADS15*/*16* and *TgMADS9* as low-temperature-responsive SVP-like genes involved in floral induction (Lu et al., 2023), reinforcing the conserved role of this gene family in temperature-mediated flowering regulation across geophytic species. SVP homologs in Lilium and *Narcissus tazetta* have also been reported as flowering repressors, though their temperature responsiveness remains unexplored (Li et al., 2015; Wu et al., 2024). Notably, ectopic expression of *CsatSVP2* in Arabidopsis caused floral morphology defects, reminiscent of phenotypes observed with Lilium SVP expression (Wu et al., 2024), suggesting that these homologs may also contribute to flower development in addition to regulating flowering time.

## Conclusion

Collectively, our findings establish a detailed molecular framework for temperature-regulated flowering in saffron, centered on the dynamic balance between floral activators (CsatFT3, CsatFD2) and repressors (CsatTFL1-3, CsatSVP2). The spatiotemporal expression patterns and protein interactions among these components provide key insights into how flowering is precisely modulated in response to environmental temperature fluctuations. By uncovering saffron-specific adaptations within otherwise conserved gene networks, this study advances our understanding of floral regulation in geophytes and sterile crops. Importantly, these results lay a foundation for the development of genetic and biotechnological strategies to improve saffron yield, synchronize flowering, and enhance resilience to climate variability. Our work highlights a saffron-specific model of thermoresponsive flowering that builds upon conserved floral regulatory pathways, yet exhibits functional divergence shaped by the geophytic lifecycle. Beyond saffron, these findings offer a valuable framework for translational research in other temperature-sensitive geophytes such as tulip, Narcissus, Hyacinthus, and Iris, where molecular insights remain scarce (Khodorova and Boitel-Conti, 2013). By identifying novel regulatory targets, this study opens new avenues for improving flowering and productivity in saffron and related high-value geophytic species.

## Acknowledgments

We gratefully acknowledge the financial support from the Council of Scientific and Industrial Research (CSIR) under the Genome Editing Mission for Crop Improvement (MMP25301), the intramural CSIR grant to IHBT (MLP201), the SERB-Start-up Research Grant (SRG/GAP0288), and the Department of Biotechnology (DBT/GAP0307) to Rajesh Kumar Singh. Senior Research Fellowships provided to Diksha Kalia (UGC), Joel Jose-Santhi (CSIR), and Firdous Rasool Sheikh (CSIR) are also gratefully acknowledged. We sincerely thank Dr. Rishikesh Bhalerao, Umeå Plant Science Centre, Swedish University of Agricultural Sciences, Sweden, for his careful reading of the manuscript and for providing constructive feedback, which significantly improved the clarity and quality of the work. Our sincere thanks go to Dr. Rimpy Diman, Technical Officer at CSIR-IHBT, for her valuable assistance with confocal microscopy. This manuscript represents the CSIR-IHBT communication number “5819”.

## Author contribution statement

R.K.S. conceptualized the research idea and designed the work plan. D.K., J.J.S., and F.R.S. conducted the experiments and collected the data. D.K., J.J.S., and R.K.S. analyzed the data and contributed to writing the manuscript. All authors read and approved the final version of the manuscript.

## Conflicts of Interest

The authors declare no conflicts of interest.

## Notes

### Competing Interest Statement

The authors have declared no competing interest.

### Summary of Updates

This version of the manuscript has been revised to incorporate minor revisions to strengthen the manuscript's scientific clarity.

## References

Abe, M., Kobayashi, Y., Yamamoto, S., Daimon, Y., Yamaguchi, A., Ikeda, Y., Ichinoki, H., Notaguchi, M., Goto, K. and Araki, T. (2005) FD, a bZIP protein mediating signals from the floral pathway integrator FT at the shoot apex. Science 309, 1052–1056.

Ahrazem, O., Rubio-Moraga, A., Nebauer, S.G., Molina, R.V. and Gomez-Gomez, L. (2015) Saffron: Its Phytochemistry, Developmental Processes, and Biotechnological Prospects. J Agric Food Chem 63, 8751–8764.

Andrés, F. and Coupland, G. (2012) The genetic basis of flowering responses to seasonal cues. Nature Reviews Genetics 13, 627–639.

Balasubramanian, S., Sureshkumar, S., Lempe, J. and Weigel, D. (2006) Potent induction of Arabidopsis thaliana flowering by elevated growth temperature. PLoS genetics 2, e106.

Bellinazzo, F., Manders, I., Heidemann, B., Bolanos, M.A., Stouten, E., Busscher, J., Abarca, D., van der Wal, F., Dornelas, M.C. and Angenent, G.C. (2025) Differential growth and flowering capacity of tulip bulbs and the potential involvement of PHOSPHATIDYLETHANOLAMINE-BINDING PROTEINS (PEBPs). Biology Direct 20, 29.

Blázquez, M.A., Trénor, M. and Weigel, D. (2002) Independent control of gibberellin biosynthesis and flowering time by the circadian clock in Arabidopsis. Plant Physiology 130, 1770–1775.

Bowman, J.L., Smyth, D.R. and Meyerowitz, E.M. (2012) The ABC model of flower development: then and now. Development 139, 4095–4098.

Capovilla, G., Schmid, M. and Posé, D. (2015) Control of flowering by ambient temperature. Journal of Experimental Botany 66, 59–69.

Cardone, L., Castronuovo, D., Perniola, M., Cicco, N. and Candido, V. (2020) Saffron (Crocus sativus L.), the king of spices: An overview. Scientia Horticulturae 272, 109560.

Freytes, S.N., Canelo, M. and Cerdán, P.D. (2021) Regulation of flowering time: when and where? Current Opinion in Plant Biology 63, 102049.

Gu, X., Le, C., Wang, Y., Li, Z., Jiang, D., Wang, Y. and He, Y. (2013) Arabidopsis FLC clade members form flowering-repressor complexes coordinating responses to endogenous and environmental cues. Nature Communications 4, 1947.

Heide, O.M. (1977) Photoperiod and temperature interactions in growth and flowering of strawberry. Physiologia Plantarum 40, 21–26.

Hsu, C.-Y., Adams, J.P., Kim, H., No, K., Ma, C., Strauss, S.H., Drnevich, J., Vandervelde, L., Ellis, J.D. and Rice, B.M. (2011) FLOWERING LOCUS T duplication coordinates reproductive and vegetative growth in perennial poplar. Proceedings of the National Academy of Sciences 108, 10756–10761.

Hu, J., Liu, Y., Tang, X., Rao, H., Ren, C., Chen, J., Wu, Q., Jiang, Y., Geng, F. and Pei, J. (2020) Transcriptome profiling of the flowering transition in saffron (Crocus sativus L.). Scientific Reports 10, 9680.

Jefferson, R.A., Kavanagh, T.A. and Bevan, M.W. (1987) GUS fusions: beta-glucuronidase as a sensitive and versatile gene fusion marker in higher plants. Embo j 6, 3901–3907.

Jin, S. and Ahn, J.H. (2021) Regulation of flowering time by ambient temperature: repressing the repressors and activating the activators. New Phytologist 230, 938–942.

Jin, S., Nasim, Z., Susila, H. and Ahn, J.H. (2021) Evolution and functional diversification of FLOWERING LOCUS T/TERMINAL FLOWER 1 family genes in plants. Seminars in Cell & Developmental Biology 109, 20–30.

Jing, S., Jiang, P., Sun, X., Yu, L., Wang, E., Qin, J., Zhang, F., Prat, S. and Song, B. (2023) Long-distance control of potato storage organ formation by SELF PRUNING 3D and FLOWERING LOCUS T-like 1. Plant communications 4.

Jose-Santhi, J., Sheikh, F.R., Kalia, D. and Singh, R.K. (2023) Sugar metabolism mediates temperature-dependent flowering induction in saffron (Crocus sativus L.). Environmental and Experimental Botany 206, 105150.

Kalia, D., Jose-Santhi, J., Kumar, R. and Singh, R.K. (2022) Analysis of PEBP Genes in Saffron Identifies a Flowering Locus T Homologue Involved in Flowering Regulation. Journal of Plant Growth Regulation, 1–20.

Kalia, D., Jose-Santhi, J., Kumar, R. and Singh, R.K. (2023) Analysis of PEBP genes in saffron identifies a flowering locus T homologue involved in flowering regulation. Journal of Plant Growth Regulation 42, 2486–2505.

Kalia, D., Jose-Santhi, J., Sheikh, F.R., Singh, D. and Singh, R.K. (2024) Tobacco rattle virus-based virus-induced gene silencing (VIGS) as an aid for functional genomics in Saffron (Crocus sativus L.). Physiol Mol Biol Plants 30, 749–755.

Khodorova, N.V. and Boitel-Conti, M. (2013) The role of temperature in the growth and flowering of geophytes. Plants 2, 699–711.

Khosa, J., Bellinazzo, F., Kamenetsky Goldstein, R., Macknight, R. and Immink, R.G.H. (2021) PHOSPHATIDYLETHANOLAMINE-BINDING PROTEINS: the conductors of dual reproduction in plants with vegetative storage organs. Journal of Experimental Botany 72, 2845–2856.

Kinoshita, A. and Richter, R. (2020) Genetic and molecular basis of floral induction in Arabidopsis thaliana. Journal of Experimental Botany 71, 2490–2504.

Komiya, R., Ikegami, A., Tamaki, S., Yokoi, S. and Shimamoto, K. (2008) Hd3a and RFT1 are essential for flowering in rice.

Lee, C., Kim, S.J., Jin, S., Susila, H., Youn, G., Nasim, Z., Alavilli, H., Chung, K.S., Yoo, S.J. and Ahn, J.H. (2019) Genetic interactions reveal the antagonistic roles of FT/TSF and TFL1 in the determination of inflorescence meristem identity in Arabidopsis. The Plant Journal 99, 452–464.

Lee, J.H., Ryu, H.-S., Chung, K.S., Posé, D., Kim, S., Schmid, M. and Ahn, J.H. (2013a) Regulation of Temperature-Responsive Flowering by MADS-Box Transcription Factor Repressors. Science 342, 628–632.

Lee, J.H., Yoo, S.J., Park, S.H., Hwang, I., Lee, J.S. and Ahn, J.H. (2007) Role of SVP in the control of flowering time by ambient temperature in Arabidopsis. Genes & development 21, 397–402.

Lee, R., Baldwin, S., Kenel, F., McCallum, J. and Macknight, R. (2013b) FLOWERING LOCUS T genes control onion bulb formation and flowering. Nat Commun 4, 2884.

Leeggangers, H.A., Nijveen, H., Bigas, J.N., Hilhorst, H.W. and Immink, R.G. (2017a) Molecular Regulation of Temperature-Dependent Floral Induction in Tulipa gesneriana. Plant Physiol 173, 1904–1919.

Leeggangers, H.A., Nijveen, H., Bigas, J.N., Hilhorst, H.W. and Immink, R.G. (2017b) Molecular regulation of temperature-dependent floral induction in Tulipa gesneriana. Plant physiology 173, 1904–1919.

Li, X.-F., Wu, W.-T., Zhang, X.-P., Qiu, Y., Zhang, W., Li, R., Xu, J., Sun, Y., Wang, Y. and Xu, L. (2015) Narcissus tazetta SVP-like gene NSVP1 affects flower development in Arabidopsis. Journal of Plant Physiology 173, 89–96.

Lu, J., Qu, L., Xing, G., Liu, Z., Lu, X. and Han, X. (2023) Genome-Wide Identification and Expression Analysis of the MADS Gene Family in Tulips (Tulipa gesneriana). Genes (Basel) 14.

Molina, R., Valero, M., Navarro, Y., Guardiola, J. and Garcia-Luis, A. (2005a) Temperature effects on flower formation in saffron (Crocus sativus L.). Scientia horticulturae 103, 361–379.

Molina, R.V., M., V., Y., N., A., G.-L. and and Guardiola, J.L. (2005b) Low temperature storage of corms extends the flowering season of saffron (Crocus sativus L.). The Journal of Horticultural Science and Biotechnology 80, 319–326.

Nakagawa, M., Shimamoto, K. and Kyozuka, J. (2002) Overexpression of RCN1 and RCN2, rice TERMINAL FLOWER 1/CENTRORADIALIS homologs, confers delay of phase transition and altered panicle morphology in rice. The Plant Journal 29, 743–750.

Nakano, Y., Higuchi, Y., Sumitomo, K. and Hisamatsu, T. (2013) Flowering retardation by high temperature in chrysanthemums: involvement of FLOWERING LOCUS T-like 3 gene repression. Journal of Experimental Botany 64, 909–920.

Navarro, C., Abelenda, J.A., Cruz-Oró, E., Cuéllar, C.A., Tamaki, S., Silva, J., Shimamoto, K. and Prat, S. (2011) Control of flowering and storage organ formation in potato by FLOWERING LOCUS T. Nature 478, 119–122.

Noy-Porat, T., Cohen, D., Mathew, D., Eshel, A., Kamenetsky, R. and Flaishman, M.A. (2013) Turned on by heat: differential expression of FT and LFY-like genes in Narcissus tazetta during floral transition. Journal of experimental botany 64, 3273–3284.

Oda, A., Narumi, T., Li, T., Kando, T., Higuchi, Y., Sumitomo, K., Fukai, S. and Hisamatsu, T. (2012) CsFTL3, a chrysanthemum FLOWERING LOCUS T-like gene, is a key regulator of photoperiodic flowering in chrysanthemums. Journal of Experimental Botany 63, 1461–1477.

Pin, P.A., Benlloch, R., Bonnet, D., Wremerth-Weich, E., Kraft, T., Gielen, J.J. and Nilsson, O. (2010) An antagonistic pair of FT homologs mediates the control of flowering time in sugar beet. Science 330, 1397–1400.

Putterill, J., Laurie, R. and Macknight, R. (2004) It’s time to flower: the genetic control of flowering time. BioEssays 26, 363–373.

Pyo, Y., Park, S., Xi, Y. and Sung, S. (2014) Regulation of flowering by vernalisation in Arabidopsis. In: Advances in botanical research pp. 29-61. Elsevier.

Renau-Morata, B., Nebauer, S.G., García-Carpintero, V., Cañizares, J., Gómez Minguet, E., de los Mozos, M. and Molina, R.V. (2021) Flower induction and development in saffron: Timing and hormone signalling pathways. Industrial Crops and Products 164, 113370.

Simon, R., Igeño, M.I. and Coupland, G. (1996) Activation of floral meristem identity genes in Arabidopsis. Nature 384, 59–62.

Singh, D., Sharma, S., Jose-Santhi, J., Kalia, D. and Singh, R.K. (2023) Hormones regulate the flowering process in saffron differently depending on the developmental stage. Front Plant Sci 14, 1107172.

Srikanth, A. and Schmid, M. (2011) Regulation of flowering time: all roads lead to Rome. Cellular and molecular life sciences 68, 2013–2037.

Strasser, B., Alvarez, M.J., Califano, A. and Cerdán, P.D. (2009) A complementary role for ELF3 and TFL1 in the regulation of flowering time by ambient temperature. The Plant Journal 58, 629–640.

Sureshkumar, S., Dent, C., Seleznev, A., Tasset, C. and Balasubramanian, S. (2016) Nonsense-mediated mRNA decay modulates FLM-dependent thermosensory flowering response in Arabidopsis. Nat Plants 2, 16055.

Tamaki, S., Matsuo, S., Wong, H.L., Yokoi, S. and Shimamoto, K. (2007) Hd3a protein is a mobile flowering signal in rice. Science 316, 1033–1036.

Taoka, K.-i., Ohki, I., Tsuji, H., Kojima, C. and Shimamoto, K. (2013) Structure and function of florigen and the receptor complex. Trends in plant science 18, 287–294.

Tsaftaris, A., Pasentsis, K. and Argiriou, A. (2013) Cloning and Characterization of FLOWERING LOCUS T-Like Genes from the Perennial Geophyte Saffron Crocus (Crocus sativus). Plant Mol. Biol. Report. 31, 1558–1568.

Tsuji, H., Nakamura, H., Taoka, K. and Shimamoto, K. (2013) Functional diversification of FD transcription factors in rice, components of florigen activation complexes. Plant Cell Physiol 54, 385–397.

Turck, F., Fornara, F. and Coupland, G. (2008) Regulation and identity of florigen: FLOWERING LOCUS T moves center stage. Annu. Rev. Plant Biol. 59, 573–594.

Tylewicz, S., Tsuji, H., Miskolczi, P., Petterle, A., Azeez, A., Jonsson, K., Shimamoto, K. and Bhalerao, R.P. (2015) Dual role of tree florigen activation complex component FD in photoperiodic growth control and adaptive response pathways. Proceedings of the National Academy of Sciences 112, 3140–3145.

Wang, Z., Li, X., Xu, J., Yang, Z. and Zhang, Y. (2021) Effects of ambient temperature on flower initiation and flowering in saffron (Crocus sativus L.). Scientia Horticulturae 279, 109859.

Wickland, D.P. and Hanzawa, Y. (2015) The FLOWERING LOCUS T/TERMINAL FLOWER 1 gene family: functional evolution and molecular mechanisms. Molecular plant 8, 983–997.

Wigge, Philip A. (2011) FT, A Mobile Developmental Signal in Plants. Current Biology 21, R374–R378.

Wigge, P.A., Kim, M.C., Jaeger, K.E., Busch, W., Schmid, M., Lohmann, J.U. and Weigel, D. (2005) Integration of spatial and temporal information during floral induction in Arabidopsis. Science 309, 1056–1059.

Wu, X., Ling, W., Pan, Y., Yang, Z., Ma, J., Yang, Y., Xiang, W., Zhou, L., Sun, M., Chen, J., Chen, H., Zheng, S., Zeng, J. and Li, Y. (2024) Functional analysis of a lily SHORT VEGETATIVE PHASE ortholog in flowering transition and floral development. Plant Physiology and Biochemistry 206, 108287.

Wu, Y.-M., Ma, Y.-J., Wang, M., Zhou, H., Gan, Z.-M., Zeng, R.-F., Ye, L.-X., Zhou, J.-J., Zhang, J.-Z. and Hu, C.-G. (2022) Mobility of FLOWERING LOCUS T protein as a systemic signal in trifoliate orange and its low accumulation in grafted juvenile scions. Horticulture Research 9.

Xue, W., Shi, J., Li, Z., Zhang, Y., Tang, M., Du, X., An, Z., Chen, H., Yang, D. and Li, X. (2025) CmFDa-mediated epigenetic regulation of flowering in chrysanthemum. Plant Biotechnology Journal.

Yan, X., Cao, Q.-Z., He, H.-B., Wang, L.-J. and Jia, G.-X. (2021) Functional analysis and expression patterns of members of the FLOWERING LOCUS T (FT) gene family in Lilium. Plant Physiology and Biochemistry 163, 250–260.

Zheng, R., Meng, X., Hu, Q., Yang, B., Cui, G., Li, Y., Zhang, S., Zhang, Y., Ma, X. and Song, X. (2023) OsFTL12, a member of FT-like family, modulates the heading date and plant architecture by florigen repression complex in rice. Plant Biotechnology Journal 21, 1343–1360.

Zhu, Y., Klasfeld, S., Jeong, C.W., Jin, R., Goto, K., Yamaguchi, N. and Wagner, D. (2020) TERMINAL FLOWER 1-FD complex target genes and competition with FLOWERING LOCUS T. Nature communications 11, 5118.

Zik, M. and Irish, V.F. (2003) Global identification of target genes regulated by APETALA3 and PISTILLATA floral homeotic gene action. The Plant Cell 15, 207–222.

