## Supplementary_Information for "Florigen Activation Complex Dynamics and SVP-Mediated Repression Orchestrate Temperature-Regulated Flowering in Saffron"

**Supplementary Figure 1**

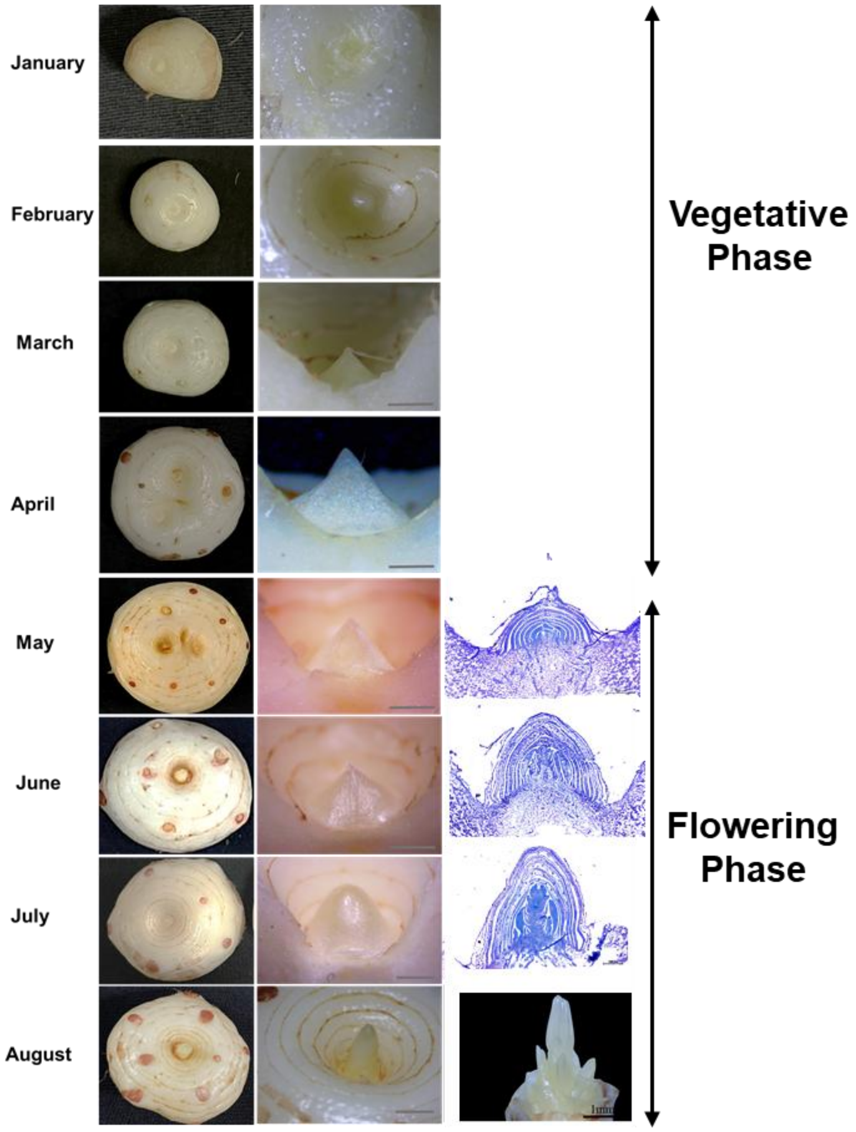


**Supplementary Figure 1. Morphological Stages of Apical Bud Differentiation Leading to Flowering in Saffron.** Representative images showing the morphological and developmental transitions during apical bud outgrowth in saffron, culminating in flowering. Saffron corms were maintained under controlled conditions as described in the Methods section. Floral induction is observed in June, followed by progressive flower differentiation and development. *Scale bar = 1 mm.*

**Supplementary Figure 2**

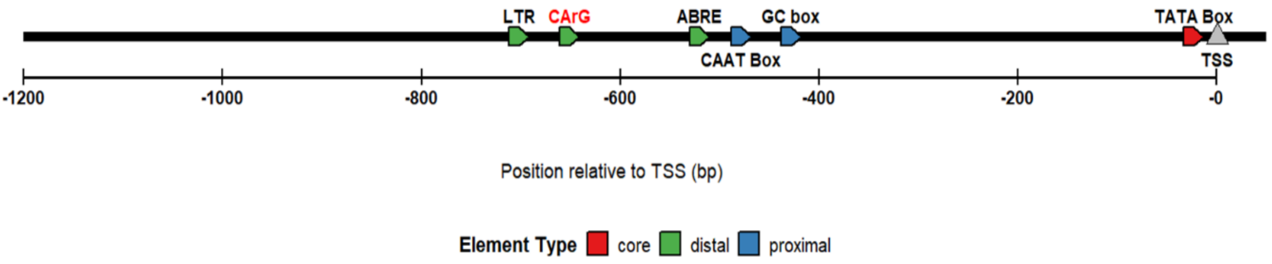


**Supplementary Figure 2: Analysis of cis-regulatory elements in the promoter regions of *CsatFT3* gene.** Schematic diagram representing the core, distal, and proximal promoter elements of CsatFT3. Distribution and positioning of cis-elements within the 1.2 kb promoter regions of CsatFT3 gene. Different types of cis-elements are marked along the promoter region. Colors and shapes represent distinct cis-element types (e.g., TATA box, MYB-binding site, etc.).

**Supplementary Figure 3**

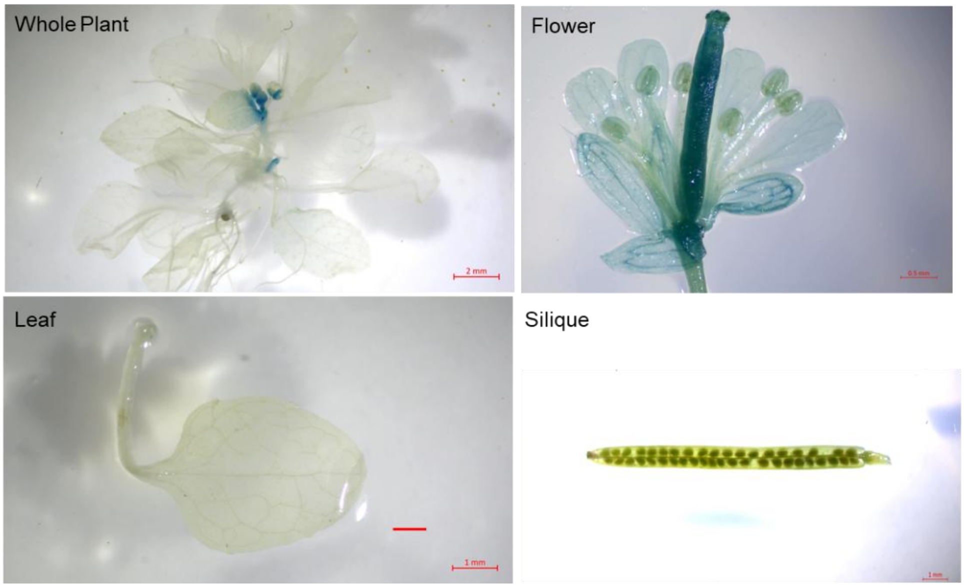


**Supplementary Figure 3: GUS expression patterns driven by the *CsatFT3* promoter in transgenic *Arabidopsis thaliana*.** Histochemical GUS staining highlights promoter activity in various tissues, including the whole plant, flower, leaf, and silique, demonstrating the spatial expression profile of the *CsatFT3* promoter**.**

### Supplementary Figure 4

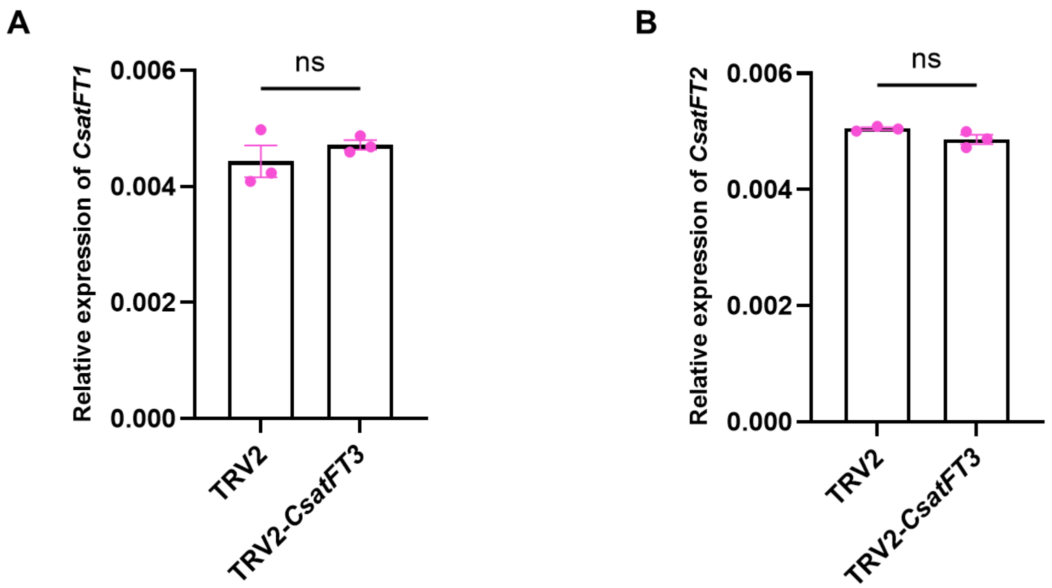


**Supplementary Figure 4. Relative expression of (A) *CsatFT1* and (B) *CsatFT2* in apical bud tissues of *CsatFT3* VIGS-treated plants, quantified by RT–qPCR.** Expression levels were normalized to a reference gene and are presented relative to mock treated controls. Values represent the mean ± standard error of mean (SEM) from three independent biological replicates. Statistical significance was analyzed using Student's t-test (*: $p<0.05$; **: $p<0.01$)

**Supplementary Figure 5**

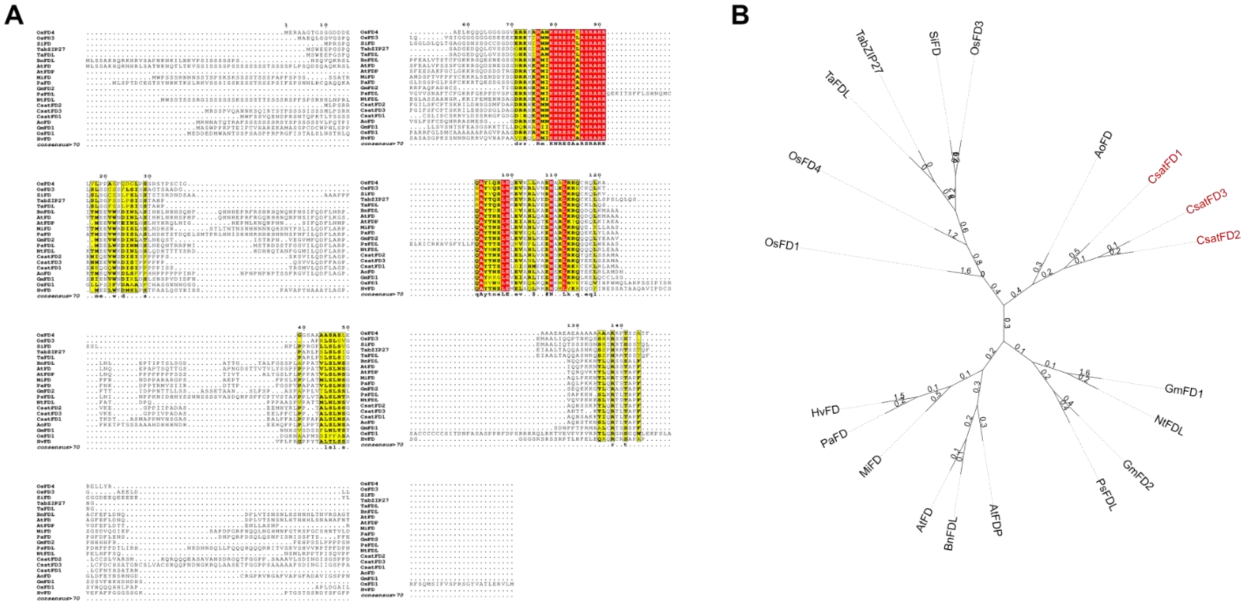


**Supplementary Figure 5: Multiple sequence alignment and phylogenetic analysis of *CsatFD* genes. (A)** Multiple Sequence Alignment of *CsatFD* Genes **(B)** Phylogenetic tree of *CsatFD* Genes. *GmFD1* (NP_001236963.1), *GmFD2* (XP_006573542.2), *OsFD1* (Os09g0540800), *OsFD3* (Os02g0833600), *OsFD4* (Os08g0549600), *PsFDL* (XP_050907953.1), *SiFD* (XP_004961973.1), *AoFD* *(*XP_020262433.1), *MiFD* (XP_044510751.1), *PaFD* (XP_034889738.1), *BnFDL* (XP_048607215.1), *NtFDL* (XP_016476358.1), *TabZIP27* (XP_044320573.1), *TaFDL* (XP_044320574.1), *AtFD* (AT4G35900), *AtFDP* (AT2G17770). Values are Branch Length

**Supplementary Figure 6**

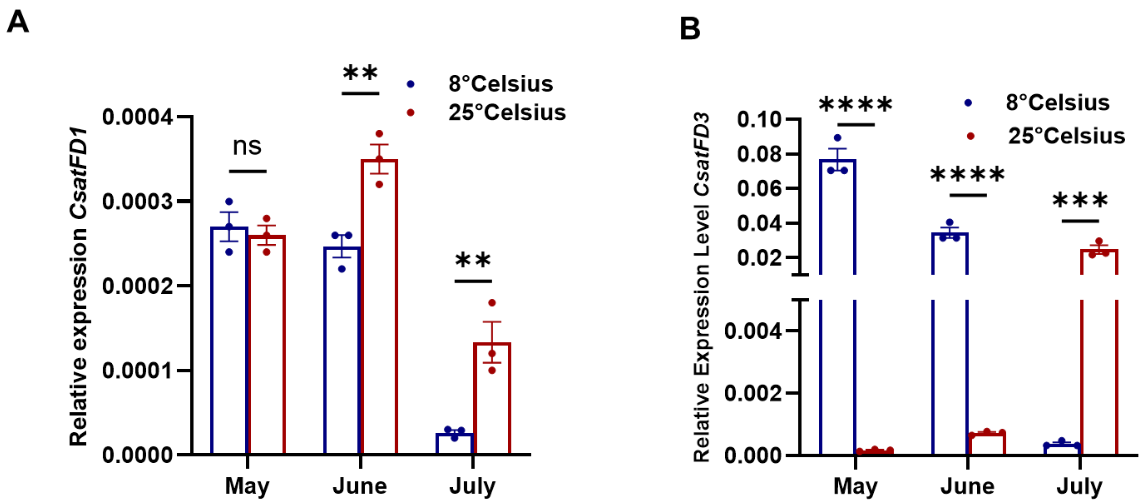


**Supplementary Figure 6: Expression patterns of CsatFD1 and CsatFD3 under different temperature conditions. (A)** *CsatFD1* and **(B)** *CsatFD3* transcript levels in apical buds collected in May, June, and July under two distinct temperature conditions were measured using RT–qPCR. Data are presented as the mean ± standard error of mean (SEM) from three independent biological replicates. Statistically significance analyzed using two-way ANOVA to assess the effects of temperature, month, and their interaction, followed by Sidak's multiple comparisons test. Asterisks indicate significant differences between 8°C and 25°C within each month (**** $p < 0.0001$)

**Supplementary Figure 7**

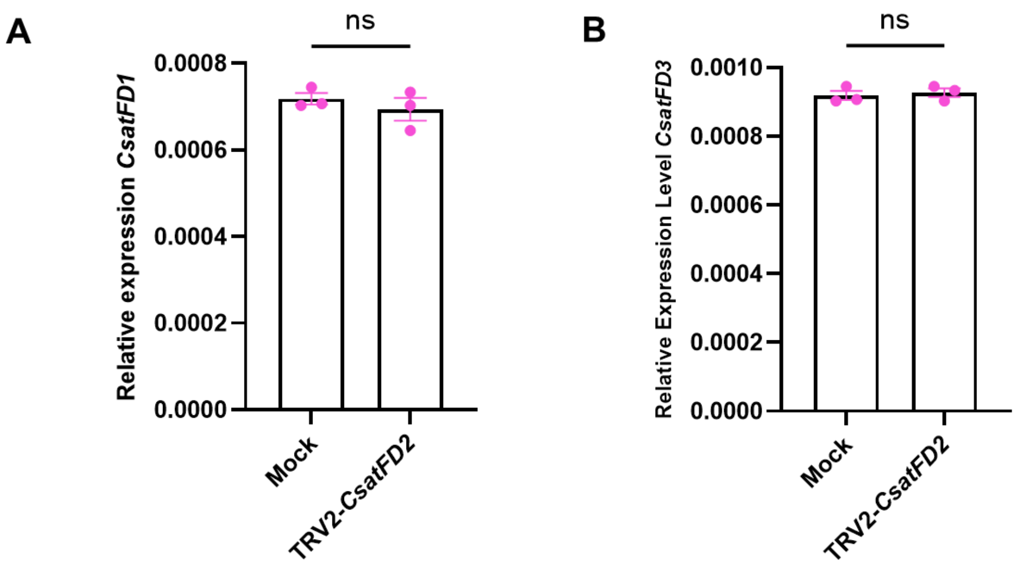


**Supplementary Figure 7: Quantitative expression analysis of *CsatFD1* and CsatFD3 in CsatFD2-silenced lines during floral induction. (A)** *CsatFD1* and **(B)** *CsatFD3* transcript levels were measured in apical buds of TRV2–CsatFD2–silenced plants using RT–qPCR. Expression values were normalized against a reference gene and calculated relative to untreated controls. Data represent the mean ± standard error of mean (SEM) from three biological replicates, each consisting of 10 individual plants (n = 10). Statistical significance was analyzed using Student's t-test (*: p<0.05; **: p<0.01)

**Supplementary Figure 8**

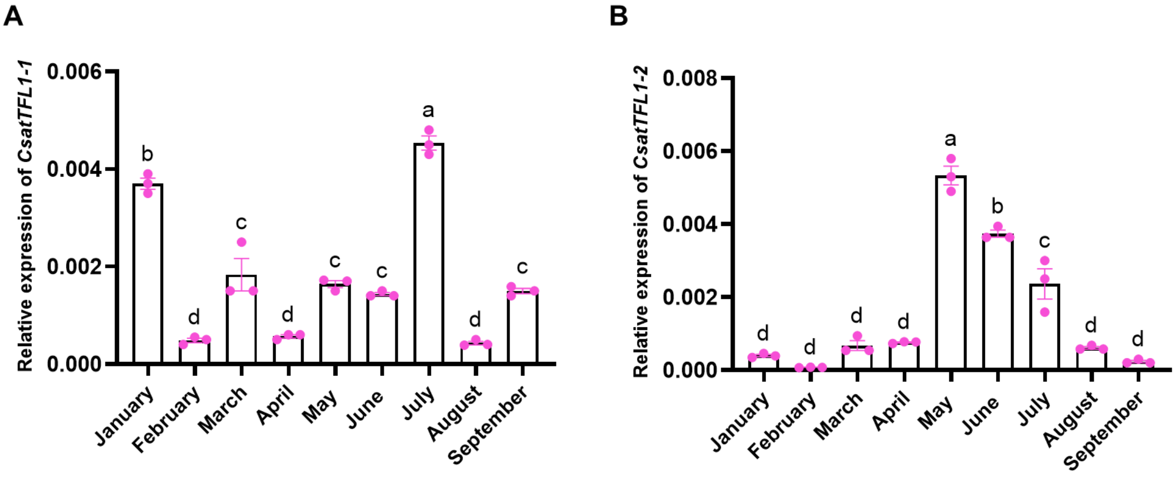


**Supplementary Figure 8: Expression profiling of *CsatTFL1*-1 and *CsatTFL1*-3 during developmental phase transitions.** (A) Relative expression of *CsatTFL1*-1 and **(B)** *CsatTFL1*-3 in apical bud tissues during the vegetative and flowering stages. Transcript levels were quantified by RT–qPCR, normalized to a reference gene, and are presented relative to the vegetative phase. Data represent the mean ± standard error of mean (SEM) from three independent biological replicates. Statistically significant differences between stages are analyzed using one-way ANOVA followed by Tukey's Honest Significant Difference (HSD) test for multiple comparisons. Different letters indicate statistically significant differences between groups at $p < 0.05$, according to Compact Letter Display (CLD).

**Supplementary Figure 9**

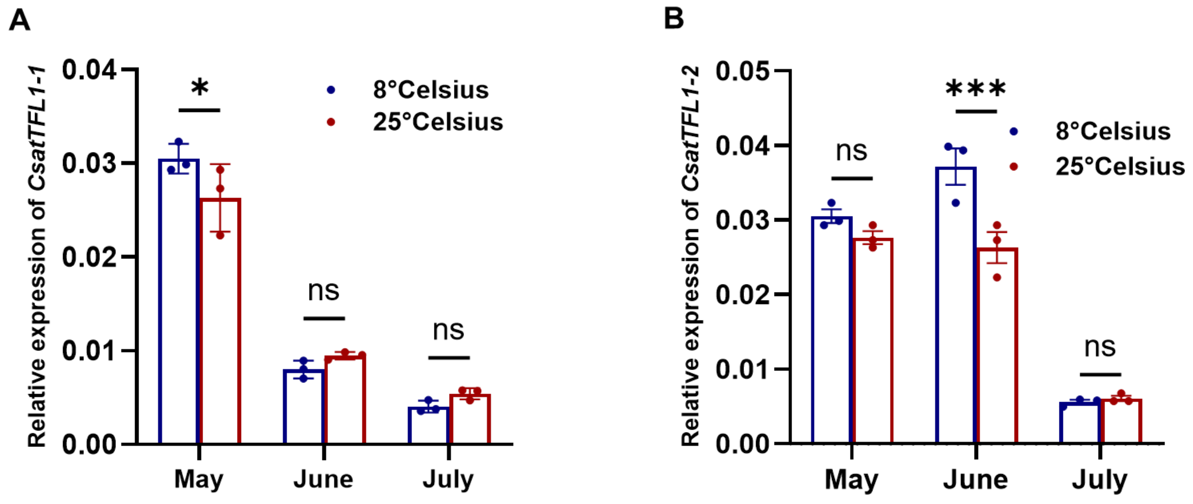


**Supplementary Figure 9: Expression of *CsatTFL1*-1 and *CsatTFL1*-2 under two temperature conditions. (A)** *CsatTFL1*-1 and **(B)** *CsatTFL1*-2 transcript levels in apical bud tissues collected in May, June, and July were assessed using RT–qPCR. The expression levels were normalized to a reference gene and presented as relative values. Data are expressed as the mean ± standard error of mean (SEM) from three independent biological replicates. Statistically significance analyzed using two-way ANOVA to assess the effects of temperature, month, and their interaction, followed by Sidak's multiple comparisons test. Asterisks indicate significant differences between 8°C and 25°C within each month (**** $p < 0.0001$)

**Supplementary Figure 10**

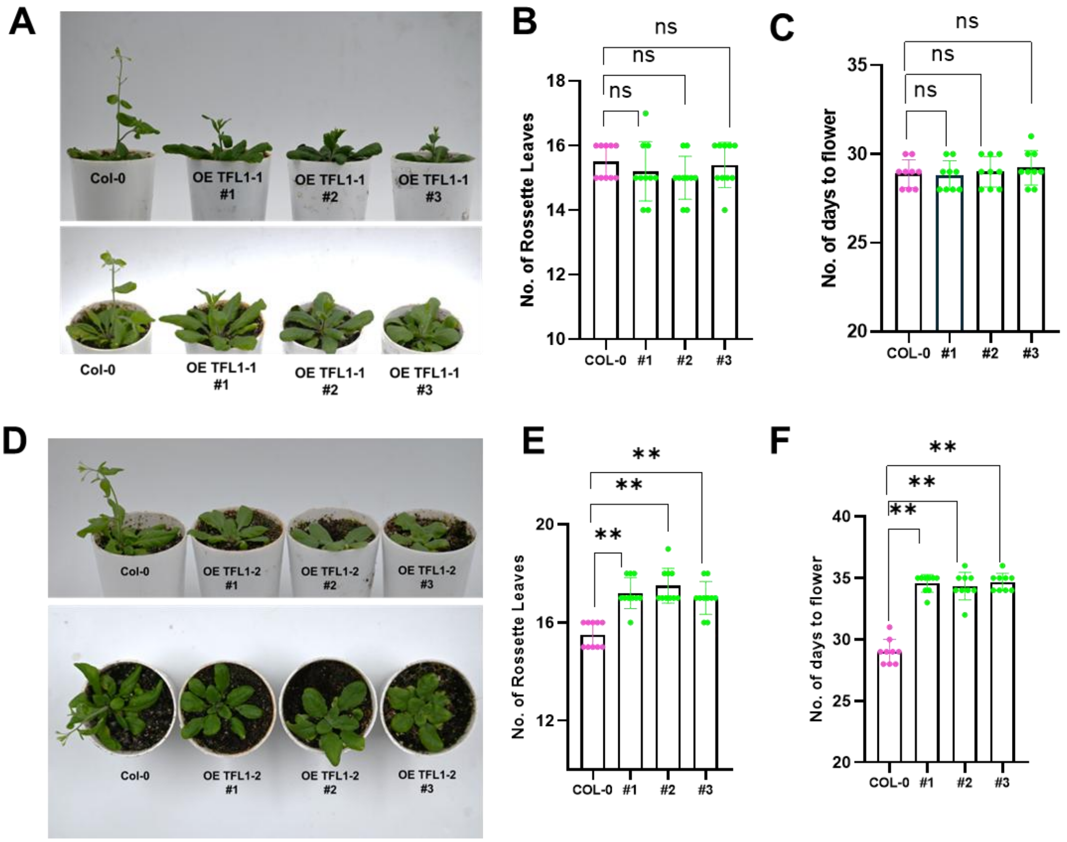


**Supplementary Figure 10: Functional analysis of *CsatTFL1-1* and *CsatTFL1-2* in Arabidopsis thaliana.** (A/D) Flowering phenotypes of 35S::CsatTFL1-1 and 35S::CsatTFL1-2 transgenic Arabidopsis lines.(B/C/E/F) Rosette leaf number and flowering time in 35S::CsatTFL1-1 and 35S::CsatTFL1-2 transgenic Arabidopsis lines. Values represent the mean ± standard error (SE) from 10 plants per replicate (n = 10). Statistical significance was analyzed using Student’s t-test (*: p<0.05; **: p<0.01)

**Supplementary Figure 11**

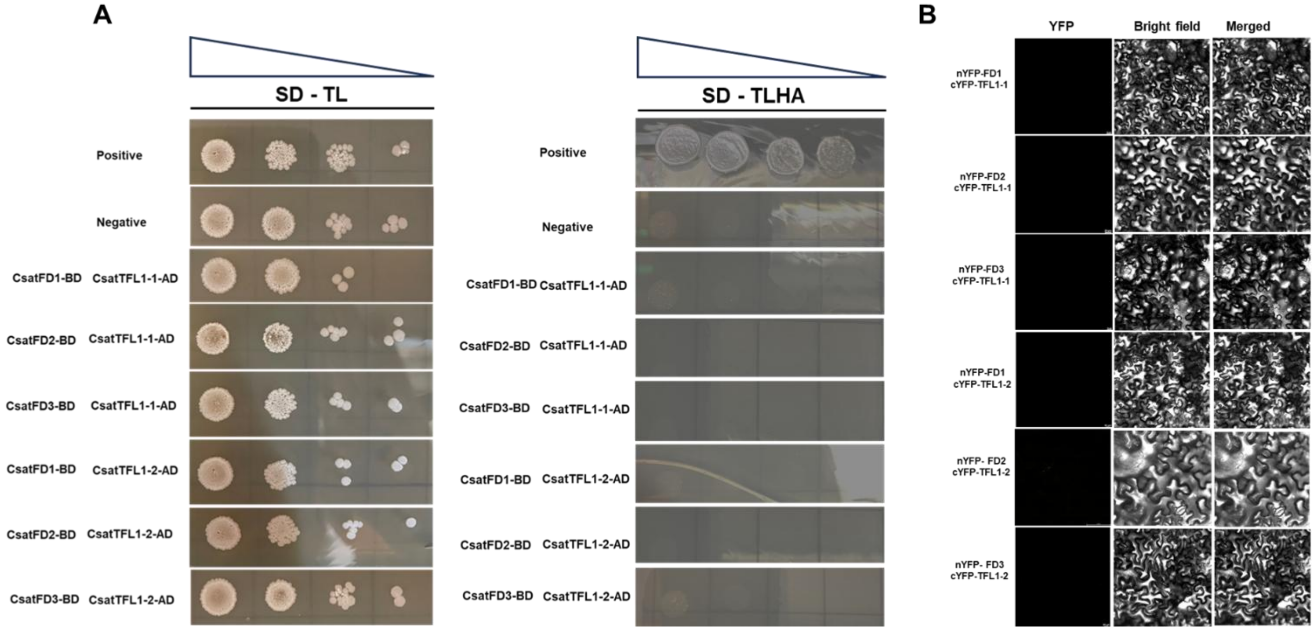


**Supplementary Figure 11: Interaction analysis of CsatTFL1-1 and CsatTFL1-2 with FD proteins. (A, B)** Yeast two-hybrid (Y2H) assay results showing no interaction between CsatTFL1-1 or CsatTFL1-2 and various FD proteins. Interactions were assessed on double dropout (DDO) SD medium lacking Trp and Leu, and quadruple dropout (QDO) SD medium lacking Trp, Leu, His, and Ade. Positive control: pGBKT7-53 and pGADT7-T; negative control: pGBKT7-lam and pGADT7-T.(C) BiFC (Bimolecular Fluorescence Complementation) assay confirms the absence of any positive interaction between CsatTFL1-1, CsatTFL1-2, and the FD proteins.

**Supplementary Figure 12**

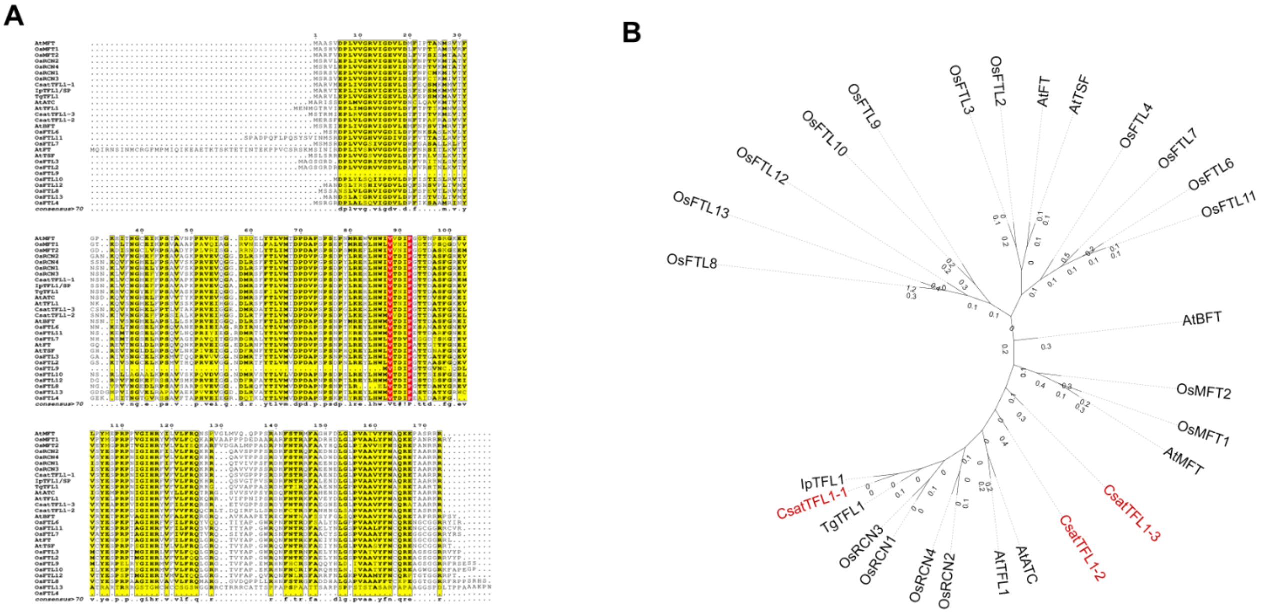


**Supplementary Figure 12: Multiple sequence alignment and phylogenetic analysis of CsatTFL1 genes. (A)** Multiple Sequence Alignment of *CsatTFL1* Genes (PEBP family) **(B)** Phylogenetic tree of *CsatTFL1* Genes. *AtATC* (CEN) (AT2G27550), *AtTFL1*(AT5G03840), *AtBFT* (Os09g0513500), *OsFTL4* (Os01g0111600), *OsMFT2* (Os01g0111600), *OsMFT1* (Os06g0498800), *AtMFT* (AT1G18100), *OsFTL13* (Os02g0232300), *OsFTL8* (Os01g0202700), *OsFTL12* (Os06g0552900), *OsFTL9* (Os01g0748800), *OsFTL10* (Os05g0518000), *OsFTL2* (Os06g0157700), *OsFTL3* (Os06g0157500), *AtTSF* (AT4G20370), *AtFT* (AT1G65480), *OsFTL7* (Os12g0232300), *OsFTL6* (Os04g0488400), *OsFTL11* (Os11g0293800), *OsRCN4* (Os04g0411400), *OsRCN2* (Os02g0531600), *OsCEN2* (Os11g0152500), *OsRCN3* (Os12g0152000), *IpTFL1* (KAJ6820878.1), *TgTFL1* (KY464932.1).

**Supplementary Figure 13**

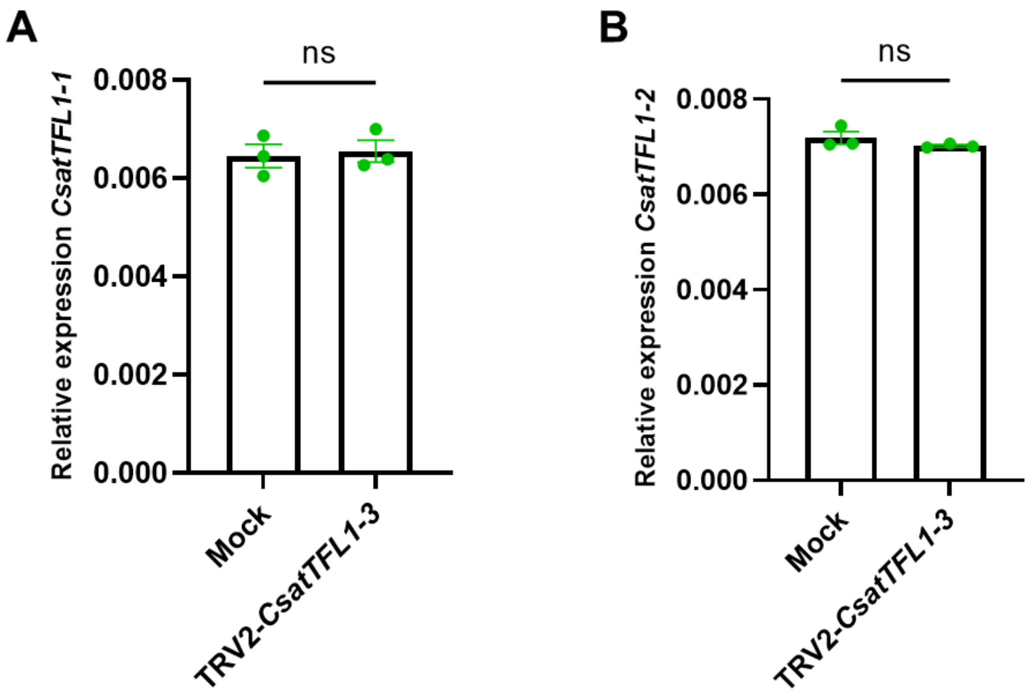


**Supplementary Figure 13: Expression of CsatTFL1-1 and CsatTFL1-2 in CsatTFL1-3 silenced lines during floral induction.** (A) CsatTFL1-1 and (B) CsatTFL1-2 transcript levels in apical bud tissues of CsatTFL1-3 silenced lines were measured by RT–qPCR. Expression levels were normalized to a reference gene. Data are presented as mean ± standard error of mean (SEM) from three independent biological replicates (n = 10). Statistical significance was analyzed using Student's t-test (*: $p<0.05$; **: $p<0.01$)

**Supplementary Figure 14**

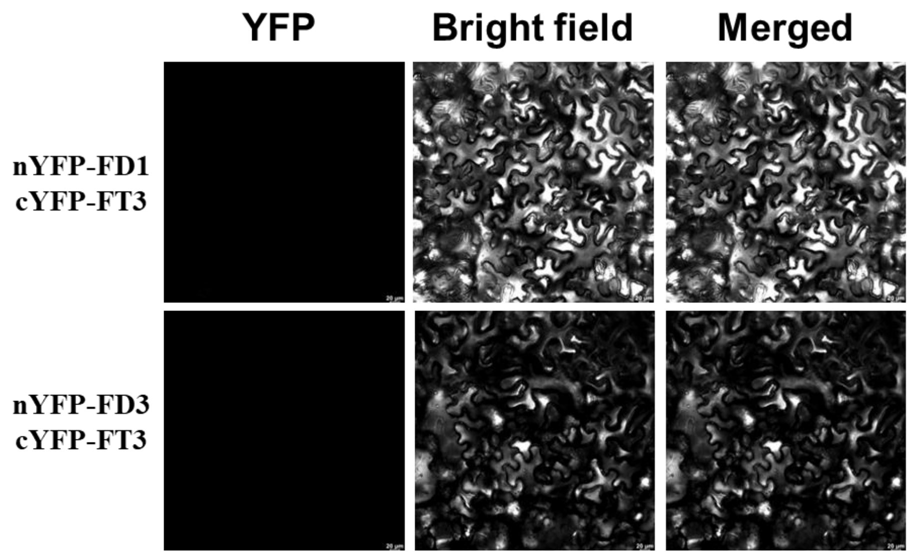


**Supplementary Figure 14: BiFC assay indicates no detectable interaction between CsatFT3 and CsatFD1 or CsatFD3.** Bimolecular Fluorescence Complementation (BiFC) was used to assess potential protein–protein interactions between CsatFT3 and CsatFD1 and CsatFD3. No reconstituted fluorescence signal was observed in Nicotiana benthamiana leaf epidermal cells co-expressing the respective BiFC constructs, suggesting that CsatFT3 does not interact with either CsatFD1 or CsatFD3.

### Supplementary Figure 15

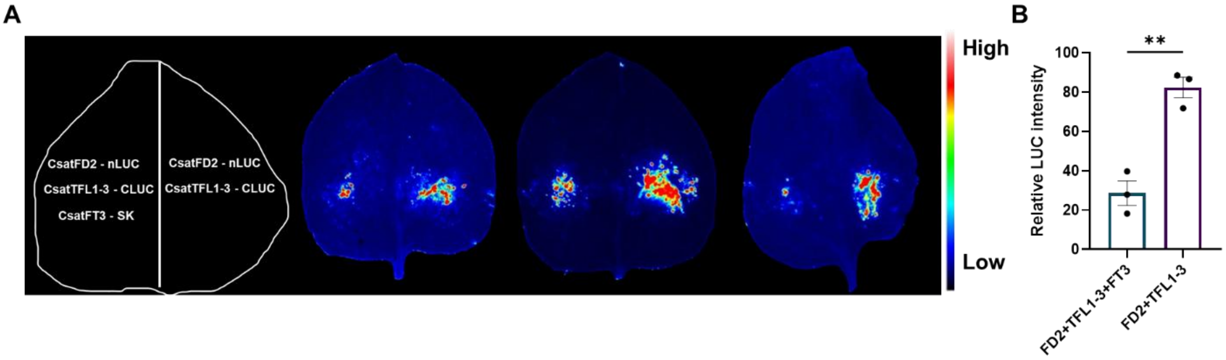


**Supplementary Figure 15: Protein–protein interaction analysis in *Nicotiana benthamiana*. (A)** LUC complementation imaging assays were used to assess the interaction between CsatTFL1-3 and CsatFD2. The C- and N-terminal halves of luciferase (LUC) were fused to CsatTFL1-3 and CsatFD2, respectively. Luminescence signals were captured 48 hours after infiltration. **(B)** The panel shows the quantified relative luciferase activity. Data were obtained from at least three independent biological replicates. Statistical significance was determined using Student's t-test: **, P < 0.01.

**Supplementary Figure 16**

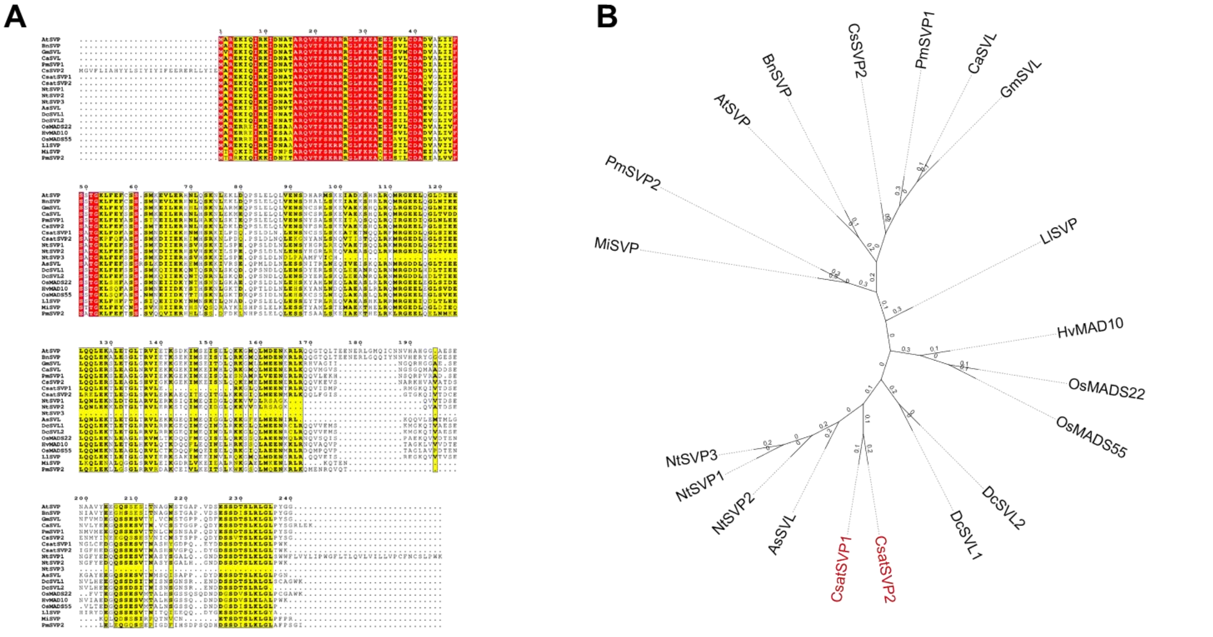


**Supplementary Figure 16: Multiple sequence alignment and phylogenetic analysis of *CsatSVP* genes. (A)** Multiple Sequence Alignment of *CsatSVP* Genes (MADS BOX family) **(B)** Phylogenetic tree of *CsatSVP* Genes. *AtSVP* (AT2G22540), *OsMADS22* (Os02t0761000-01), *OsMADS55* (Os06t0217300-01), *AsSVL* (Asa0G05496.1), *NtSVP1* (AHC92621.1), *NtSVP2* (AHC92622.1), *NtSVP3* (AHC92623.1), *DcSVP1* (XP_020692709.1), *DcSVP2* (XP_020692711.1), *PmSVP2* (XP_016650557.1), *PmSVP1* (XP_008235586). *MiSVP (*XP_044488836.1). *BnSVP* (NP_001303013.1). *HvMADS10* (XP_044952618.1). *LlSVP* (APY23912.1) *CaSVL* (XP_027078661.1). *GmSVP* (NP_001240951.1). *NtSVP1* (AHC92621.1), *NtSVP2* (AHC92622.1). *AsSVL* (Asa0G05496.1).

**Supplementary Figure 17**

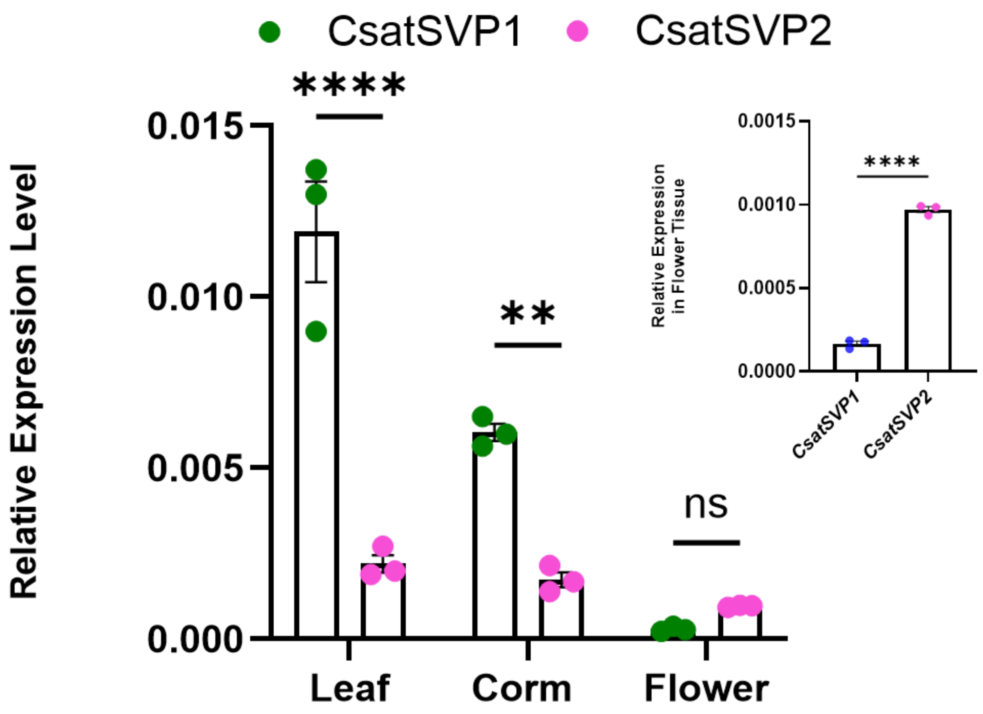


**Supplementary Figure 17: Expression profiling of *CsatSVP1* and *CsatSVP2* in different plant tissues** (Leaf, Corm, Flower). Statistically significance analyzed using two-way ANOVA to assess the effects of temperature, month, and their interaction, followed by Sidak's multiple comparisons test. Asterisks indicate significant differences between 8°C and 25°C within each month (**** $p < 0.0001$) and Student's t-test: **, $P < 0.01$.

**Supplementary Figure 18**

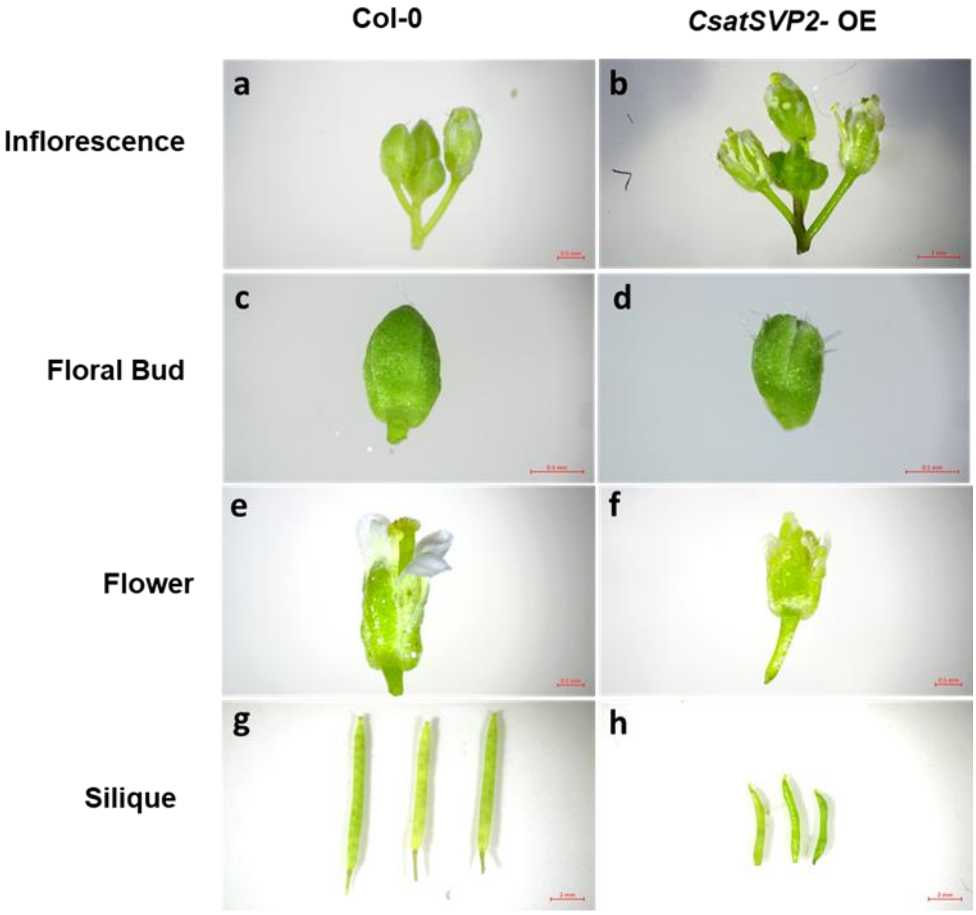


**Supplementary Figure 18: Phenotypic changes in floral organs of 35S::CsatSVP2 transgenic Arabidopsis plants.** The left panel shows the flowering phenotype of wild-type Arabidopsis (Col-0), and the right panel compares the flowering phenotype of the 35S::CsatSVP2 transgenic plants.

### Supplementary tables

### Supplementary Table 1. List of primers used in the study

| Name | Sequence of Primer (5' - 3') | Purpose |
|---|---|---|
| CsatFT3F0 | CACCATGGCCGCAGCTACTAGAGAAACGG | Overexpression studies |
| CsatFT3R0 | AGCGTACGTTCTCCTCCTCCCACCAGAGCC | |
| Csat FD1 F0 | CACCATGTGGTTCTCTTCGGTACAAGAGA | |
| CsatFD1 R0 | TCAAAATGGAGCTGTTAAGGTTCTCTGG | |
| CsatFD2 F0 | CACCATGCTTCCTTCTAGAAGGAGGAACATTC | |
| CsatFD2 R0 | TCAAAATGGAGCTGTTAAGGTTCTCTGG | |
| CsatFD3F0 | CACCATGCGGTCCTCACCAGTGCAGGCG | |
| CsatFD3R0 | GCGAACATTAACAGCTCCATTT | |
| CsatTFL1-1F0 | CACCATGACTAGAATGATAGACCCAC | |
| CsatTFL1-1R0 | TCATCTTCTTCTTGCAGCGGT | |
| CsatTFL1-2 F0 | CACCATGGCAGAGTACATAAAAGACAC | |
| CsatTFL1-2R0 | CTATAAAAACCGTCCACCCACTGA | |
| CsatTFL1-3F0 | CACCATGACTAGAATGATAGAGCCAC | |
| CsatTFL1-3R0 | TCATCTTCTTCTTGCAGCGGT | |
| CsatSVP1 F0 | CACCATGGCGAGGGAGAAGATACAAATAAGG | |
| CsatSVP1 R0 | TTACTTCCATGGAAGACCTAACTTG | |
| CsatSVP2 F0 | CACCATGGCGAGGGAGAAGATACAGATAAG | |
| CsatSVP2 R0 | GTTTATTCTGCTTACTTCCATGTAAGC | |
| CSFT3VIGSF | CGGAATTCATGGCCGCAGCTACTAGAGAAACGG | VIGS Studies |
| CsFT3VIGSR | GAATTCTCCGGTGCTCCCAGGAATATCTA | |
| Cs FD2 ViGS F0 | CGGGATCCATGCTTCCTTCTAGAAGGAGG | |
| CsFD2 ViGS R0 | CGGGATCCCCAAAAGTCTGCCCCCTGTC | |
| CsCEN2VigsF | GCGAATTCATGACTAGAATGATAGAGCC | |
| CsCEN2VigsR | GCGGATCCAATGTGGCATTAGTGGTGCC | |
| CsSVP2 ViGS F0 | CGGAATTCATGCGGAGGGAGGAAGAT | |
| CsSVP2 ViGS R0 | CGGGATCCCTGCTGAGTTGAGATCGTT | |
| CsFT3ProR2 | GCCATATGGAGACCCTGGCCTTATAAG | Luciferase Assay |
| CsatFT3PRO T1LA | GCGGTACCCCTGATGGATTCACCGAAAC | |
| CsatFT3PRO T2LA | GCGGTACCGGACTTACCGTTTGATTGAT | |
| CsTubulinF | TGATTTCCAACTCGACCAGTGTC | Real Time PCR |
| CsTubulinR | ATACTCATCACCCTCGTCACCATC | |
| CsFT3Fq | ATATGGCCGCAGCTACTAGAGAAAC | |
| CsFT3Rq | TCCACCATCACCAGAGTGTAGAAAT | |
| CsFD1Fq | ATGTGGTTCTCTTCGGTACAAGAGA | |
| CsFD1Rq | CCATAGATGCCGTATCTTTTGTGGG | |
| CsFD2Fq | ATGCTTCCTTCTAGAAGGAGGAACATTC | |
| CsFD2Rq | TCAGCGGGAATAATCGGTGGGCCCTCC | |
| CsatFD3Fq | ATGCGGTCCTCACCAGTGCAGGCG | |

| | | |
|---|---|---|
| CsatFD3Rq | TGGCCGAGTGGCAGTCGAAGCAGAG | |
| CsTFL1-1 Fq | GGCAGAAGCGCCGTCAGTC | |
| Cs TFL1-1 Rq | AGCGGCGTGTACTGAACCGATCC | |
| Cs TFL1-2 Fq | CCGCCTCACCGATCAGAT | |
| Cs TFL1 -2 Rq | CGAAACTTCGGGTGCAGAAG | |
| CsatTFL1-3Fq | ATGACTAGAATGATAGAGCCAC | |
| CsatTFL1-3Rq | AGTGACAATCCAGTGAAGGTGT | |
| SVP Fq | ATCCCTCAAGTTAGGTCTTCCAT | |
| SVP Rq | GTCACAAGTTTTTCTCACCCAACT | |
| SVP2 Fq | ACAACTCTTCGGTATTTCCGGAACC | |
| SVP2 Rq | ACCCAACTCGTTCTGTTTATTCTGC | |
| CsFT3ProR2 | GCCATATGGAGACCCTGGCCTTATAAG | Yeast one hybrid |
| CsatFTProY1HF0 | GCGAGCTCCGACGGCCCGGGCTTGTAAA | |
| CsatFTPROY1HR0 | GCCCCGGGATGGAGACCCTGGCCTTATA | |
| CsatSVP2Y1HF0 | GCGAATTCATGGCGAGGGAGAAGATACA | |
| CsatSVP2Y1HR0 | GCGGATCCTTACTTCCATGTAAGCCCT | |
| CsatTFL1-3 FFLC F0 | GCGGTACCATGACTAGAATGATAGAGCC | Firefly complementation assay |
| CsatTFL1-3FFLC R0 | GCGGATCCTCATCTTCTTCTTGCAGCGGT | |
| CsatFD2 FFLC F0 | GCGGTACCATGCTTCCTTCTAGAAGGAGG | |
| CsatFD2FFLCR0 | GCGGATCCAAATGGAGCTGTTAAGGTTC | |
| CsatFT3FFLAF0 | GCGAATTCATGGCCGCAGCTACTAGAGAA | |
| CsatFT3FFLAR0 | GCAAGCTTTCAAGCGTACGTTCTCCTCC | |
| CaMV35SF | GTAAGGGATGACGCACAATCC | Vector Primer |
| GUS-R | ATTCCACAGTTTTCGCGATCCAGAC | |
| AP2 | ACTATAGGGCACGCGTGGT | |
| pGADT7-F0 | CTATTCGATGATGAAGATACCCCACCAAACC | |
| pGADT7-R0 | GTGAACTTGCGGGGTTTTTCAGTATCTACGATT | |
| M13 Forward | GTA AAA CGA CGG CCA GTG | |
| M13 Forward | GGT TTT CCC AGT CAC GAC | |
| TRV2F | ATGAGCTTTATTATTACGGACGAGTG | |
| TRV2R | TTCAGACACGGATCTACTTAAAGAAC | |
| CsatFD1 R0 | AAATGGAGCTGTTAAGGTTCTCTGG | BIFC Assay |
| CsatFD2R0 | AAATGGAGCTGTTAAGGTTCTCTGG | |
| CsatFD3R0 | GCGAACATTAACAGCTCCATTT | |
